# Proteomic analysis uncovers clusterin-mediated disruption of actin-based contractile machinery in the trabecular meshwork to lower intraocular pressure

**DOI:** 10.1101/2024.02.16.580757

**Authors:** Avinash Soundararajan, Ting Wang, Padmanabhan P Pattabiraman

**Affiliations:** Ophthalmology, Indiana University School of Medicine, Indianapolis, Indiana, United States; Medical Neuroscience Graduate Program, Stark Neuroscience Research Institute, Indiana University School of Medicine, Indianapolis, Indiana, USA

**Keywords:** Clusterin, intraocular pressure, global proteomics, actin-cytoskeleton, extracellular matrix

## Abstract

Glaucoma, a major cause of blindness, is characterized by elevated intraocular pressure (IOP) due to improper drainage of aqueous humor via the trabecular meshwork (TM) outflow pathway. Our recent work identified that loss of clusterin resulted in elevated IOP. This study delves deeper to elucidate the role of clusterin in IOP regulation. Employing an *ex vivo* human anterior segment perfusion model, we established that constitutive expression and secretion as well as exogenous addition of clusterin can significantly lower IOP. Interestingly, clusterin significantly lowered transforming growth factor β2 (TGFβ2)-induced IOP elevation. This effect was linked to the suppression of extracellular matrix (ECM) deposition and, highlighting the crucial role of clusterin in maintaining ECM equilibrium. A comprehensive global proteomic approach revealed the broad impact of clusterin on TM cell structure and function by identifying alterations in protein expression related to cytoskeletal organization, protein processing, and cellular mechanics, following clusterin induction. These findings underscore the beneficial modulation of TM cell structure and functionality by clusterin. Specifically, clusterin influences the actin-cytoskeleton and focal adhesion dynamics, which are instrumental in cell contractility and adhesion processes. Additionally, it suppresses the activity of proteins critical in TGFβ2, G-protein, and JAK-STAT signaling pathways, which are vital for the regulation of ocular pressure. By delineating these targeted effects of clusterin within the TM outflow pathway, our findings pave the way for novel treatment strategies aimed at mitigating the progression of ocular hypertension and glaucoma through targeted molecular interventions.

## 1. Introduction

Glaucoma is a major global health concern, and one of the leading causes of irreversible blindness, posing a significant public health challenge [1]. This complex optic neuropathy is primarily defined by the progressive degeneration of retinal ganglion cells, leading to significant optic nerve damage [2]. Among the various types of glaucoma, primary open-angle glaucoma (POAG) is notably prevalent, characterized by increased intraocular pressure (IOP) and optic nerve deterioration [3, 4]. Understanding the intricate mechanisms behind IOP regulation is crucial, particularly highlighting the role of the trabecular meshwork (TM) outflow pathway as a key component in this regulatory system. The TM outflow pathway comprises the specialized contractile TM cells and the extracellular matrix (ECM) alongside juxtacanalicular tissue (JCT) and Schlemm’s canal (SC) in the regulation of aqueous humor (AH) drainage [5].

The TM tissue is constantly subjected to several physiological and pathological stressors [6], which are countered by the adaptive responses of the tissue to achieve homeostasis. However, in our previous study, we showed that the levels of a stress response chaperone protein clusterin were decreased under several IOP dysregulators in TM [7] resulting in a loss of clusterin function. Further, we found that functional loss of clusterin produces a significant increase in IOP [7] which was independently validated by another study [8]. Noticeably, our study uncovered that at the cellular level, reduced clusterin triggered significant actin reorganization evidenced by increased levels of actin-associated proteins and membrane proteins like integrins[7]. Additionally, there was a marked rearrangement of focal adhesions such as paxillin and vinculin along stress fibers. Previous studies have shown that the build-up of the ECM and the reorganization of the actin-cytoskeleton, led to changes in the stiffness and contractility of the TM, culminating in elevated resistance to aqueous humor drainage [9–12]. Dysregulated canonical transforming growth factor β2 (TGFβ2) signaling involving mothers against decapentaplegic (SMADs) as well as the activation of Rho GTPase play a major role in regulating actin contractility and ECM changes [11, 13–15]. These changes are mediated by the pro-fibrotic response in the TM tissue, leading to increased resistance to AH drainage [16–18]. Among the signaling cascades, the activation of protein kinase C (PKC), PI3K-Akt pathway, and mitogen-activated protein kinases (MAPKs) pathway influence actin dynamics, crucial for the structural and functional integrity of the TM [19–22]. This also involves altering the gene expression related to actin dynamics, actin filament stabilization, and polymerization [20, 23, 24].

Interestingly, our previous study using a targeted biochemical approach demonstrated that constitutive induction of clusterin secretion in TM mitigated actin-related cytoskeletal polymerization machinery, ECM, and pro-fibrotic proteins [7]. Thus, suggesting the potential of clusterin in IOP lowering. This study aimed to explore the functional and mechanistic roles of clusterin in maintaining IOP homeostasis using anterior segment perfusion cultures *ex vivo*. By employing a comprehensive global proteomic approach, we sought to identify the complex molecular interactions and pathways influenced by clusterin. This will serve to uncover the key signaling pathways and molecular networks vital for IOP regulation, offering insights for diagnosis and treatment paradigms of ocular hypertension and POAG.

## 2. Methodology

### 2.1 Materials

The reagents and antibodies used in this study are presented in **Table 1**.

**Table 1.** Materials.

| **Reagents and antibodies** | **Catalog number** | **Company** |
| --- | --- | --- |
| Recombinant human clusterin | CLUH5227100UG | AcroBiosystems |
| Recombinant TGFβ2 | GF440 | Millipore Sigma |
| Anti-collagen 1 | 600-401-103 | Rockland |
| Anti-clusterin | sc6420 | Santa Cruz Biotechnology |
| Anti-paxillin | sc365379 | Santa Cruz Biotechnology |
| Anti-GAPDH | sc-25778 | Santa Cruz Biotechnology |
| Anti-Gα q/11/14 | sc-365906 | Santa Cruz Biotechnology |
| Anti-MLC2 | 3672 | Cell Signaling Technology |
| Anti-phospho-MLC2 | 3671 | Cell Signaling Technology |
| Anti-fibronectin | - | Gift from Dr. Harold Erickson, Duke University |
| Donkey-anti-goat IgG | 705-035-003 | Jackson Immuno Research |
| Donkey-anti-mouse IgG | 715-035-150 | Jackson Immuno Research |
| Donkey-anti-rabbit IgG | 715-035-144 | Jackson Immuno Research |
| Goat-anti-Mouse IgG Alexa Fluor Plus 488 | A32723 | Thermo Fisher Scientific |
| Donkey-anti-Mouse IgG Alexa Fluor 555 | A31570 | Thermo Fisher Scientific |
| Donkey-anti-Rabbit IgG Alexa Fluor Plus 488 | A32731 | Thermo Fisher Scientific |
| Phalloidin (Alexa Fluor Plus 488 PHA) | A12379 | Invitrogen |
| Fluoromount-G | E139999 | Invitrogen |
| DAPI | UH2820551 | Sigma-Aldrich |
| ViraPower Adenoviral Expression System | K4930-00 | Invitrogen |
| Advantage-HF 2 PCR kit | 639123 | Clontech |
| pENTR/D-TOPO cloning vector | K243520 | Invitrogen |
| pAd/CMV/V5-DEST vector | V493-20 | Invitrogen |
| LR Clonase II enzyme mix | 11791020 | Invitrogen |
| Pac 1 | RO547S | New England Biolabs |
| Adeno-X™ Maxi Purification | 631533 | Takara Bio |
| Antibiotic-Antimycotic | 15240062 | Gibco |
| Background Sniper | BS966H | Biocare Medicals |
| Tween 20 | TWEEN201 | MP Biomedicals |
| Fluoroshield mounting medium with DAPI | ab104139 | Abcam |
| OptiMEM | 31985-070 | Gibco |
| penicillin-streptomycin-glutamine solution | 10378-016 | Gibco |
| Collagenase type 4 | 9007-34-5 | Worthington Biochemical |
| Human serum albumin | A4329 | MilliporeSigma |
| 199 media | 11150-059 | Gibco |
| 5X All-In-One RT MasterMix | G492 | Applied Biological<br>Materials |
| 2X qPCR MasterMix-ROX | MasterMix-LR-XL | Applied Biological<br>Materials |
| Tris Triton Buffer | 42020310 | bioWORLD |
| Pierce protease and phosphatase inhibitor | A3961 | Thermo Fisher Scientific |
| Trizol LS reagent | 10296028 | Life technologies |
| HEPES | H-4034 | Sigma-Aldrich |
| Trichloroacetic acid | A322-100 | Fisher Chemical |
| Gelatin from porcine skin | G1890 | Sigma-Aldrich |
| Dithiothreitol | 20290 | Thermo Fisher Scientific |
| Sucrose | S1697 | Spectrum Chemical |
| Bromophenol Blue | B392-5 | Fisher Chemical |
| Urea | 161-0731 | Bio-Rad |
| Tris hydrochloride | 10812846001 | Sigma-Aldrich |
| 1.5 ml Bioruptor Plus TPX microtubes | C30010010-300 | Diagenode |
| Bioruptor Plus sonication device | B01020001 | Diagenode |
| Bio-Rad Protein Assay Dye Reagent | 5000006 | Bio-Rad |
| tris(2-carboxyethyl) phosphine hydrochloride | C4706 | Sigma-Aldrich |
| Trypsin/Lys-C Mix | V5072 | Promega |
| Sep-Pak® Vac cartridges | WAT054955 | Waters |
| Pierce Quantitative colorimetric assay | 23275 | Thermo Fisher Scientific |
| TMT10plex™ Isobaric Label Reagents and Kits | 90309 | Thermo Fisher Scientific |
| High Select™ Phosphopeptide Enrichment Kits &<br>Reagents | A32993 | Thermo Fisher Scientific |
| Orbitrap Exploris 480™ mass spectrometer | BRE725539 | Thermo Fisher Scientific |
| EasySpray™ C18 column | ES902A | Thermo Fisher Scientific |
| Acetonitrile | LS122500 | Thermo Fisher Scientific |

### 2.2 Construction of replication-deficient recombinant adenovirus expressing clusterin

Generation of replication-defective recombinant adenovirus constitutively expressing secretory clusterin (AdCLU) under CMV promoter was generated as previously described [7] from human clusterin in pcDNA6 (a gift from Claudia Koch-Brandt, Johannes Gutenberg University of Mainz) [25]. A control empty virus (AdEMPTY or AdMT) a gift from Dr. Douglas Rhee, Case Western Reserve University; was also amplified and purified.

### 2.3 Human anterior segment perfusion model *ex vivo*

Freshly enucleated human whole globes were obtained from VisionFirst, Indiana Lions Eye Bank, and Lions Gift of Sight, Minneapolis within 24-48 hours post retrieval. Details of donor age, sex, and race are provided in **supplementary table 1**. The eye globes were cleaned by trimming the extra-ocular muscle and quickly immersed in povidone-iodine topical antiseptic solution (Betadine), followed by washes in 1x phosphate-buffered saline (PBS), pH 7.4 solution. The globes were then cut below the limbus, and anterior segments with intact TM were prepared by removing the remnant of the vitreous humor, lens, ciliary body, and iris. The anterior segments were then mounted onto a custom-made perfusion chamber consisting of a foundational component and an O-ring. The foundational component consisted of a convex mound with two cannulas that allow for an influx of fluid and its subsequent drainage to simulate the AH pathway. The apparatus was connected to the transducer, which constantly recorded the pressure using the LabChart 8 application on a laptop. The fluid influx through the syringe was monitored using a syringe pump set at 2.5 μl/min. Initially, a baseline was set using DMEM containing 1x Antibiotic-Antimycotic. Once the baseline is set DMEM containing either AdCLU (3 x 10^9^ IFU/ml) to induce clusterin or AdMT (1.8 x 10^9^ IFU/ml) as control or 5 ng/ml TGFβ2 or 0.5 µg/ml recombinant clusterin (rhCLU) was perfused as per the experimental design. Post perfusion tissues were collected and fixed for histological analysis.

### 2.4 **Immunofluorescence staining:**

After *ex vivo* perfusion, the portion of the eye containing TM tissues was sectioned and fixed in 4% paraformaldehyde. Paraffin-embedded human TM tissue slides were prepared at the Histology Core, Indiana University, and immunolabeling was performed. Briefly, five-micron thick tissue sections were deparaffinized in fresh xylene thrice for 5 minutes each. The sections were subsequently hydrated with ethanol dilutions (100%, 95%, and 70%). To unmask the antigen epitopes, heat-induced antigen retrieval was performed using 0.1 M citrate buffer pH 6.0 (Vector Laboratories) for 20 min at 100°C. The slides were then blocked for nonspecific interactions with Background Sniper. For immunofluorescence staining, the primary antibody was used at 1:100 dilution overnight at 4°C. The slides were subsequently washed with 1×PBST containing 0.05% Tween 20 in 1X PBS, pH 7.4, and secondary antibodies were applied at 1:200 for 1 h at room temperature. After two additional washes, the coverslips were mounted with Fluoroshield mounting medium with 4′,6-diamidino-2-phenylindole (DAPI). For staining, a minimum of two slides per ocular tissue sample were utilized, with each slide containing three consecutive sections from the same eye. This was repeated across the biological replicates tested.

HTM cells were grown on 2% gelatin-coated 4-chamber glass bottom dishes until they attained 60-70% confluency. After appropriate treatment, cells were washed with 1X PBS twice, fixed in 4% paraformaldehyde for 20 min, permeabilized with 0.2% triton-x-100 in PBS buffer for 10 min, and blocked with 5% bovine serum albumin in 1X PBS for 1 h. Cells were then incubated with the respective primary antibodies overnight at 4°C. After washing three times with 1X PBST, they were incubated in Alexa fluor-conjugated secondary antibodies for 1 h at room temperature. Finally, the dishes were washed with PBST and were imaged under a Zeiss LSM 700 confocal microscope, z-stack images were obtained, and processed using Zeiss ZEN image analysis software. Fluorescence intensities were measured using Image J 1.53S.

### 2.5 **Scanning electron microscopy:**

Post-perfusion tissues were fixed immediately in Karnovsky solution containing 4% paraformaldehyde and 2.5% glutaraldehyde in PBS, pH 7.4 for 24 hours at 4°C. Tissues were then washed in 1X PBS, pH 7.4, and then post-fixed in 1% osmium tetraoxide for 2 hours at 4°C. The tissues were then dehydrated in increasing grades of acetone series (30-100%) and dried in a critical point dryer. The tissue sections were then mounted on small metal stubs and coated with colloidal gold. The specimens were examined under the JEOL7800F Field Emission Scanning Electron Microscope at Integrated Nanosystems Development Institute, Indiana University, Indianapolis. The images were acquired as a top view, surface morphology in lower electron detector mode at a magnification of 10KX with a pressure of 3.6 x 10^-4^ Pa in a vacuum. Images were analyzed for fibrillar matrix thickness using Image J 1.53S.

### 2.6 **Cell culture treatments:**

Primary human TM (HTM) cells were cultured from TM tissue isolated from the leftover donor corneal rings after they had been used for corneal transplantation at the Indiana University Clinical Service, Indianapolis, and characterized as described previously [7, 26]. HIPPA compliance guidelines were adhered to for the use of human tissues. The usage of donor tissues was exempt from the DHHS regulation and the IRB protocol (1911117637), approved by the Indiana University School of Medicine IRB review board. Donor age, race, and sex details are provided in **supplementary table 1**. The expanded population of HTM cells was characterized by the detection of dexamethasone-induced myocilin.

Primary porcine TM (PTM) cells were cultured from fresh porcine TM globes obtained from Indiana Packers, Delphi, IN, USA, which were transported on ice within 2 hours of sacrifice. The isolated TM tissue was finely chopped and enzymatically dissociated, using a solution containing 10 mg collagenase type 4, and 3 mg human serum albumin in 199 media at 37°C for 1 hour in a shaker. The cells were pelleted by centrifugation at 2500 rpm for 12 min, collected, and plated in a 2% gelatin-coated plastic tissue culture plate with DMEM containing 20% FBS and PSQ. The expanded population of PTM cells was sub-cultured in DMEM containing 10% FBS. The PTM cells were maintained at 37 °C under 5% CO2. All experiments were conducted using confluent HTM cultures or PTM as mentioned, using cells in between passages four to six. Cells were treated with AdCLU or AdMT for 72 hours with the last 48 hours in serum starved conditions. For experiments involving rhCLU (0.5 µg/ml), cells were serum-starved overnight before treatment. All experiments in this study were performed using biological replicates.

### 2.7 **Measurement of myosin light-chain (MLC) phosphorylation:**

Myosin phosphorylation analysis was performed as described previously [20, 27]. Briefly, post treatment, HTM cells were incubated with 10% cold trichloroacetic acid (TCA) for 5 min and then washed with ice-cold water five times to completely remove the TCA. The cells were extracted with 8 M urea buffer containing 20 mM Tris, 23 mM glycine, 10 mM dithiothreitol (DTT), saturated sucrose, and 0.004% Bromophenol Blue, using a sonicator. The urea-solubilized samples were separated on urea/glycerol PAGE containing 30% acrylamide, 1.5% bisacrylamide, 40% glycerol, 20 mM Tris, and 23 mM glycine. Then proteins from the glycerol gels were transferred onto nitrocellulose filters in 10 mM sodium phosphate buffer, pH 7.6. Membranes were probed using phospho-MLC (p-MLC) or MLC primary antibodies.

### 2.8 **Enrichment of ECM fraction:**

The HTM cells were treated with AdCLU for 72 hours and AdMT treatment was used as a control. Following treatment, ECM enrichment was carried out. Cells were washed twice with 1X phosphate-buffered saline (PBS) pH 7.4 and permeabilized with 0.2% Triton X-100/PBS at room temperature (RT) for 10 min. Permeabilized cells were treated with 10 ml of 0.3% ammonium hydroxide/100 cm cell culture plate and incubated for 5 min at RT. Plates were rinsed 4-5 times with ultrapure water to completely remove the cellular debris. Then enriched ECM was collected by scrapping in 25 mM HEPES, pH 7.4.

### 2.9 **Global proteomics analysis:**

#### Sample preparation for proteomics

Whole cell pellets or ECM enriched samples were lysed in 8 M Urea, 100 mM Tris-HCl, pH 8.5 by sonication in 1.5 ml Micro Tubes using a Bioruptor® Plus sonication system with 30 s/30 s on/off cycles for 15 min in a water bath at 4°C. After subsequent centrifugation at 14,000g for 20 min, protein concentrations were determined by Bradford protein assay. A 20 µg equivalent of protein from each sample was reduced with 5 mM tris(2-carboxyethyl) phosphine hydrochloride (TCEP, Sigma-Aldrich, C4706) for 30 min at room temperature, and the resulting free cysteine thiols were alkylated with 10 mM chloroacetamide for 30 min at room temperature in the dark. Samples were diluted with 50 mM Tris HCl, pH 8.5, to a final urea concentration of 2 M for Trypsin/Lys- C based overnight protein digestion at 37°C (1:100 protease: substrate ratio, mass spectrometry grade).

Peptide purification and labeling Digestions were acidified with trifluoracetic acid (TFA, 0.5% v/v) and desalted on Sep-Pak® Vac cartridges with a wash of 1 ml 0.1% TFA followed by elution in 70% acetonitrile 0.1% formic acid (FA). Peptides were dried by speed vacuum and resuspended in 29 µL of 50 mM triethylammonium bicarbonate. Peptide concentrations were checked by Pierce Quantitative colorimetric assay. The same amount of peptide from each sample was then labeled for 2 hours at room temperature with 0.2 mg of Tandem Mass Tag reagent (TMT™ Isobaric Label Reagent Set). Labeling reactions were quenched by adding 0.3% hydroxylamine (v/v) to the reaction mixtures at room temperature for 15 min. Labeled peptides were then mixed and dried by speed vacuum.

#### High pH basic fractionation

For high pH basic fractionation, peptides were reconstituted in 0.1% TFA and fractionated using methodology and reagents from Pierce™ High pH reversed-phase peptide fractionation kit.

#### Nano-LC-MS/MS analysis

Nano-LC-MS/MS analyses were performed on an EASY-nLC HPLC system coupled to Orbitrap Exploris 480™ mass spectrometer with a FAIMS pro interface. About 1/8 of each global peptide fraction and 1/4 of each phosphopeptide fraction were loaded onto a reversed-phase EasySpray™ C18 column (2 μm, 100 Å, 75 μm × 25 cm) at 400 nl/min. Peptides were eluted from 4%– 30% with mobile phase B (mobile phases A: 0.1% FA, water; B: 0.1% FA, 80% acetonitrile over 160 min, 30%–80% B over 10 min, and dropping from 80%–10% B over the final 10 min. The mass spectrometer was operated in positive ion mode with a 4 s cycle time data-dependent acquisition method with advanced peak determination. The FAIMS CV was maintained at −50 V. Precursor scans (m/z 375- 1600) were done with an orbitrap resolution of 60,000, RF lens% 40, maximum inject time 50 ms, normalized AGC target 300%, and MS2 intensity threshold of 5e4, including charges of 2–6 for fragmentation with 60 s dynamic exclusion. MS2 scans were performed with a quadrupole isolation window of 0.7 m/z, 35% HCD CE, 45,000 resolution, 200% normalized AGC target, auto maximum IT, and fixed first mass of 110 m/z.

#### Data analysis

The resulting RAW files were analyzed in Proteome Discover™ 2.4 (Thermo Fisher Scientific) with a Homo sapiens UniProt FASTA (last modified 02/15/2017) plus common contaminants. SEQUEST HT searches were conducted with a maximum number of two missed cleavages, a precursor mass tolerance of 10 ppm, and a fragment mass tolerance of 0.02 Da. Static modifications used for the search were as follows: 1) carbamidomethylation on cysteine (C) residues, 2) TMT6plex label on N-peptide N-termini, and 3) TMT6plex label on lysine (K) residues. Dynamic modifications used for the search were oxidation of methionines and acetylation, Met-loss, or Met-loss plus acetylation of protein N-termini. The percolator false discovery rate was set to a strict setting of 0.01 and a relaxed setting of 0.050. In the consensus workflow, peptides were normalized by total peptide amount with no scaling. Co-isolation of 50% and average reporter ion S/N 10 were used as thresholds for quantification. Resulting in normalized abundance values for each sample type, fold change (abundance ratio) and log2 (abundance ratio) values, and respective p-values (ANOVA) from Proteome Discover™ were exported to Microsoft Excel.

### 2.10 **Gene expression analysis:**

Gene expression analysis was carried out as mentioned in our previous study [7]. Total RNA was extracted after experiments from TM tissues and cells using the Trizol method following the manufacturer’s protocol. The RNA amounts were quantified using a NanoDrop 2000 UV-Vis Spectrophotometer (Thermo Scientific). Equal amounts of RNA were then reverse-transcribed to complementary DNA (cDNA) using the 5X All-In-One RT MasterMix with genomic DNA removal according to the manufacturer’s instructions. The following reaction condition was maintained for cDNA conversion: incubation at 25°C for 10 min, followed by incubation at 42°C for 15 min, and enzyme inactivation at 85°C for 5 min. The cDNA was diluted as per requirement, and 10 µl was used for gene expression analysis using Quant Studio Flex 6/7 thermocycler (ThermoFisher Scientific). Bright Green 2X qPCR MasterMix-ROX and gene-specific oligonucleotides (Integrated DNA Technologies) were used for the analysis. Sequence-specific forward and reverse oligonucleotide primers for the indicated genes are provided in **Table 2**.

**Table 2:**
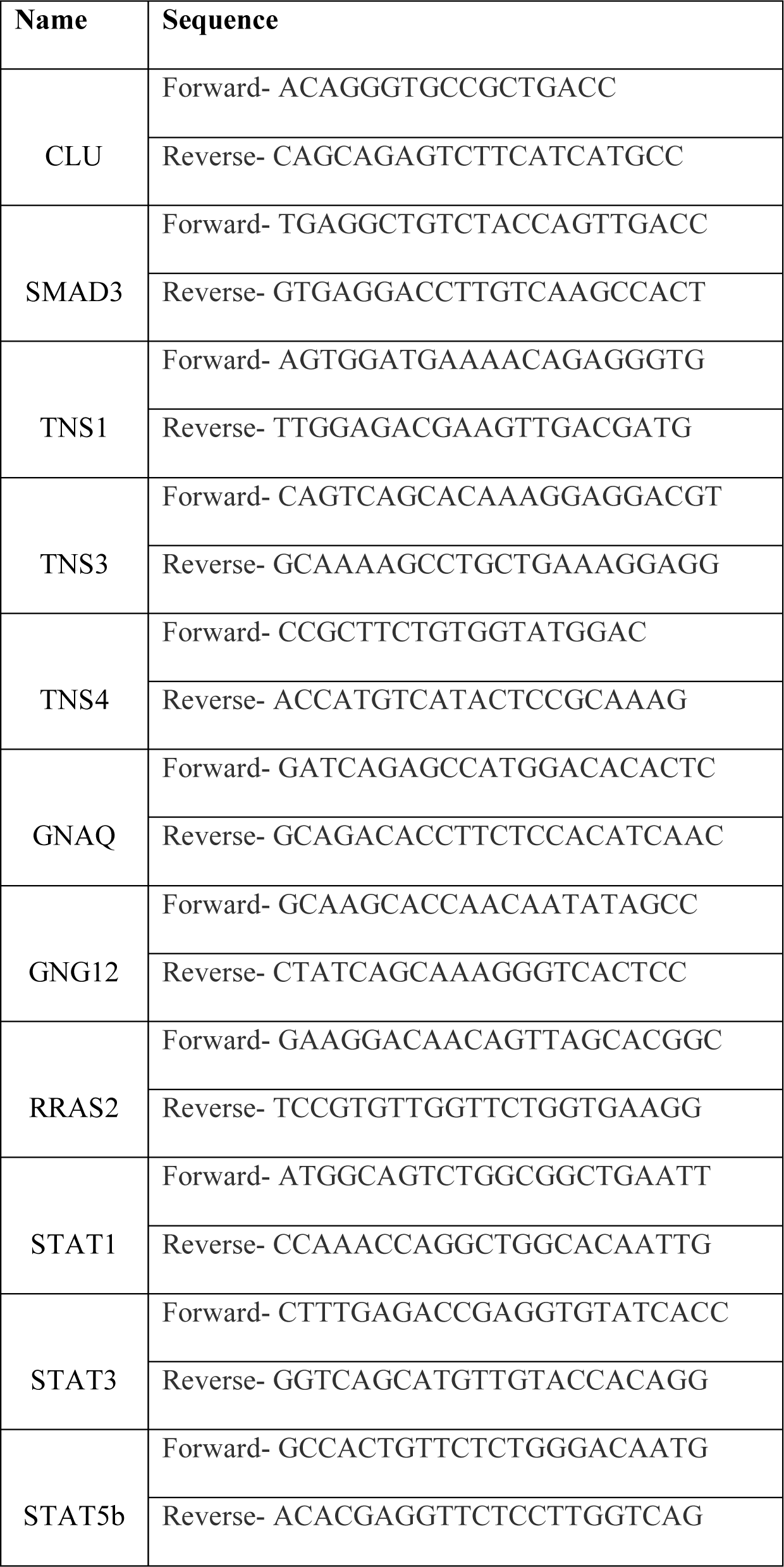
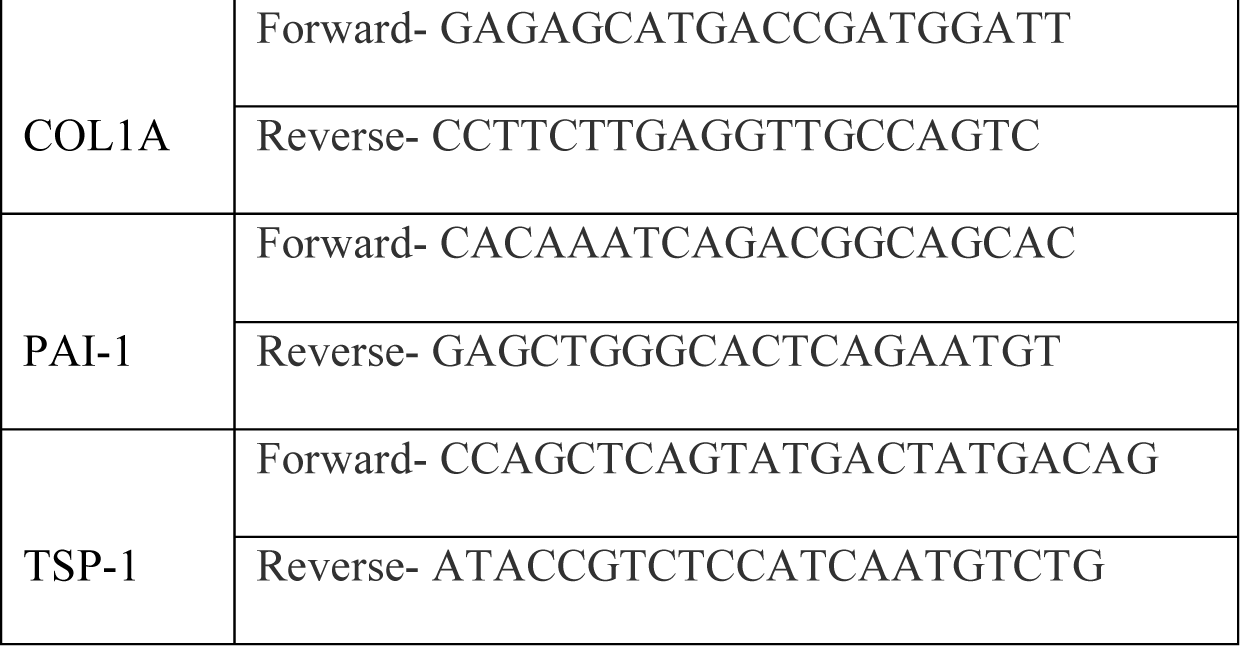
Oligonucleotide primers used in the qPCR amplification.

### 2.11 **Membrane fraction extraction:**

Membrane fraction was isolated based on previously published work with some modifications [28]. The porcine TM cells were seeded in 10 cm plates. Post treatment cells were collected by scrapping in Tris Triton Buffer containing Pierce protease and phosphatase inhibitor. Samples were then homogenized and centrifuged at 800 g for 10 min at 4^ο^C. The supernatant containing the total protein was collected and centrifuged at 40,000 rpm for 1 h at 4^ο^C using Beckman Coulter Optima MAX-XP Ultracentrifuge. The supernatant containing soluble cytoplasmic fraction was removed and stored in a fresh tube. The pellet containing the insoluble membrane fraction was washed with hypotonic buffer and centrifuged at 40,000 rpm for 1 h at 4^ο^C. The supernatant was discarded, and the pellet collected was solubilized using a urea sample buffer containing 8 M urea, 0.25 M Tris, 0.25 M Glycine, 0.5 M DTT, and saturated sucrose.

Samples were then sonicated and used for immunoblotting.

### 2.12 **Statistical analysis:**

All data presented was based on a minimum of four independent observations. The graphs represent the mean ± standard error. All statistical analyses were performed using Prism 8.1.1 (GraphPad). The difference between the two groups was assessed using paired and unpaired Student’s t-tests when parametric statistical testing was applicable. When multiple groups were compared, a one-way or two-way analysis of variance followed by the Tukey post hoc test was applied. The p-value ≤ 0.05 was considered a statistically significant difference between the test and control samples.

## 3. **Results:**

### 3.1 Constitutive clusterin expression reduced IOP in human anterior segment perfusion model (*ex vivo*) by reducing collagen 1

Using the human anterior segment perfusion model, we evaluated the effect of constitutive clusterin expression on IOP. A stable baseline IOP was established by perfusing the eyes with DMEM supplemented with antibiotic-antimycotic at the rate of 2.5 µl/min for ∼24 hours. Subsequently, AdCLU was introduced into one eye, and control AdMT virus to the contralateral eye. The eyes perfused with AdMT are denoted by a black line with blue circles, while those perfused with AdCLU are represented by a black line with red squares **(Figure 1A)**. The virus perfusion lasted for 24 hours, followed by a continuation of perfusion with DMEM for a duration of ∼336 hours (13 days). Eyes that received AdCLU showed a decrease in IOP starting from 72 hours, with this reduction being sustained over time. Notably, there was a significant decrease in IOP observed at various intervals: 192 hours (p=0.04), 216 hours (p=0.04), 312 hours (p=0.01), and 336 hours (p=0.02), across a sample size of n=5, as depicted in Figure 1A. The maximum decrease in pressure was around 23% at 192 hours with mean IOP fold change of 0.766 mmHg). The distinctive decrease in IOP observed in eyes treated with AdCLU compared to AdMT underscores the potential of clusterin to lower IOP.

**Figure 1:**
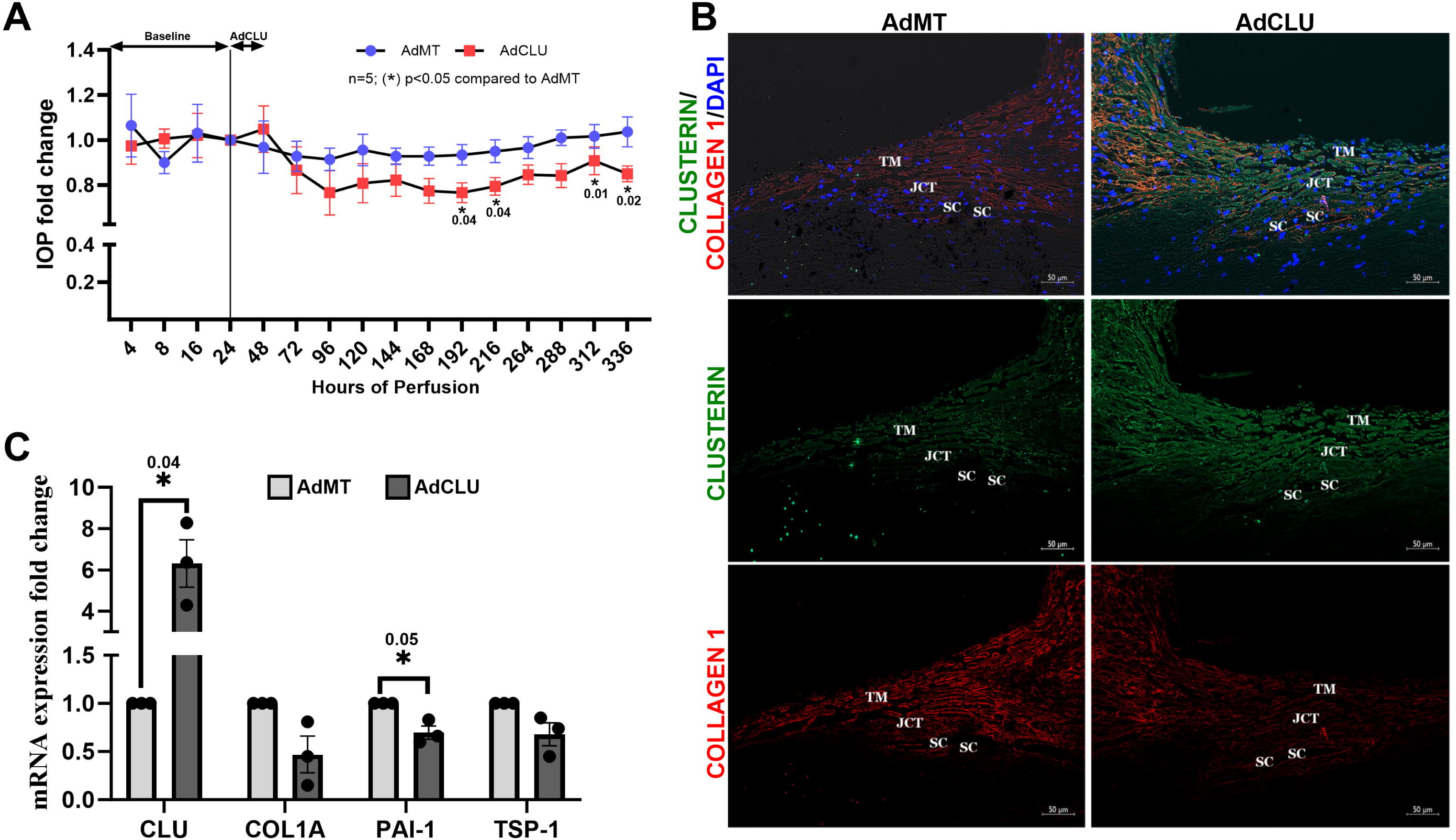
Constitutive clusterin expression lowers IOP by modifying ECM expression *ex vivo*. A) Adenovirus-mediated constitutive clusterin expression (AdCLU) significantly reduced IOP compared to control virus (AdMT). AdMT and AdCLU are represented by a black line with blue circles and a black line with red squares respectively, indicating mean IOP fold change at each given time point corresponding to the x-axis containing hours of perfusion. Error bars represent the standard error of the mean (SEM), and (*) indicates significance compared to AdMT, with a sample size of n=5. B) Immunofluorescence analysis of outflow pathway tissue shows AdCLU-induced clusterin (green color) expression and reduced collagen 1A (green color) expression compared to AdMT-perfused tissues. DAPI staining nucleus is in blue. The scale bar is 50 μm. C) Gene expression analysis shows an increase in clusterin and reduction in ECM protein collagen, pro-fibrotic PAI-1, and TSP-1 in AdCLU-perfused tissue compared to AdMT-perfused tissue. (*) indicates significance compared to AdMT, with a sample size of n=3.

After perfusion, two wedges from each of the perfused tissues were processed for histology, and the remainder of the TM tissues were used for RNA extraction. Immunofluorescence analysis in **Figure 1B** displays the comparison between tissues perfused with AdMT (left panel) and AdCLU (right panel). The top panel represents an overlay image of the distribution of clusterin (in green), Collagen 1 (in red), and nuclear staining with DAPI (in blue). As a proof of concept for successful viral transduction in the outflow pathway, tissues perfused with AdCLU exhibited a marked increase in clusterin immunopositive cells (middle right panel) within the TM-JCT region predominantly. The inner wall of SC in the outflow pathway did not light up greatly compared to the AdMT. Moreover, tissues that received AdCLU showed a marked decrease in Collagen 1 distribution (bottom right panel) across the TM outflow pathway when compared to the AdMT treated samples.

Gene expression analysis using qPCR was performed on the RNA isolated from TM tissues extracted from the perfused eyes to determine the impact of clusterin on the transcription of ECM protein COL1, the pro-fibrotic proteins plasminogen activator inhibitor 1 (PAI-1), and thrombospondin-1 (TSP1) [7]. The *GAPDH* served as the loading control. While clusterin levels were significantly elevated by ∼ 6 folds, the Pro-fibrotic PAI-1 showed significant reduction (p=0.05, n=3) and both COL1 and TSP-1 expression demonstrated a downward trend in tissues perfused with AdCLU compared to AdMT **(Figure 1C**). These findings validated our previous assessment in HTM cells *in vitro* where we demonstrated that clusterin induction decreased ECM and pro-fibrotic proteins [7].

### 3.2 Exogenous supply of clusterin reduced the pathological IOP elevation in the human anterior segment perfusion model (*ex vivo*)

Though AdCLU lowered IOP, we were unsure if this was caused due to an increase in intracellular clusterin or the secreted clusterin. To discern the functional role of the secreted clusterin on IOP regulation, we exogenously supplied clusterin to the anterior segments and recorded the changes in IOP using the human anterior segment perfusion model. After achieving stable baseline pressure, rhCLU at a concentration of 0.5 μg/ml in DMEM was perfused for approximately 145 hours (∼ 6 days). The chosen concentration for clusterin perfusion was based on insights from a previous study along with our prior research [7, 29]. Notably, there was a significant reduction in IOP (p=0.0001, n=4) starting from ∼70 hours (∼3 days) compared to the baseline, and this significant decrease was sustained consistently for ∼144 hours (∼6 days) (**Figure 2A**). This result provided a compelling rationale for the notion that secreted clusterin is significant for IOP lowering.

**Figure 2:**
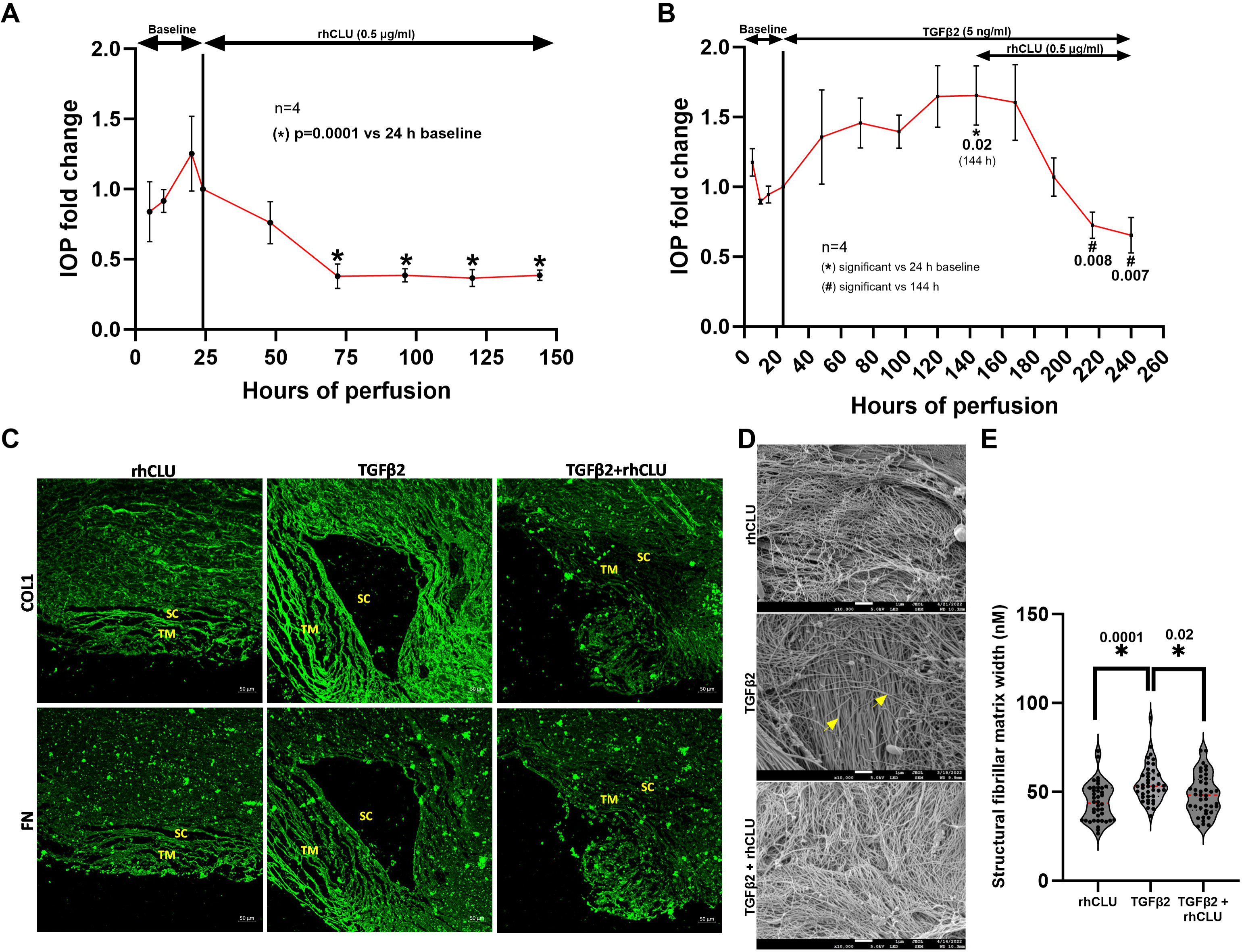
Exogenous supplementation of clusterin lowers pathological IOP elevation *ex vivo*. A) Exogenous supplementation of 0.5 μM recombinant clusterin (rhCLU) reduces IOP. Error bars represent SEM, and (**\***) indicates significance compared to 24 h of average baseline IOP, with a sample size of n=4. B) Starting from a 24-hour baseline, TGFβ2 was continuously perfused until the end of the experiment. The supplementation of rhCLU was initiated after observing an increase in IOP at 144 hours and was maintained for the duration of the perfusion. Error bars represent SEM, and (**\***) indicates significance compared to 24 h of average baseline IOP, (#) indicates significance compared to IOP elevation at 144 h, with a sample size of n=4. C) Immunofluorescence analysis shows the increase in ECM proteins-COL1A and FN in outflow pathway tissues perfused only with TGFβ2 compared to rhCLU alone. Tissues when supplemented with rhCLU along with TGFβ2 (TGFβ2 + rhCLU) reduction in COL1A and FN are seen. The scale bar is 50 μm. D) Scanning electron microscopy images showing the surface morphology of TM tissue perfused with rhCLU, TGFβ2, and TGFβ2 + rhCLU. The images were acquired in lower electron detector mode at a magnification of 10,000X with a pressure of 3.6 x 10^-4^ Pa in vacuum. Images were analyzed for fibrillar matrix thickness using Image J 1.53S. The scale bar is 1 μm. The yellow arrow in TGFβ2 shows a thickened fibrillar matrix. E) Graphical representation violin plots demonstrating the increase in structural fibrillar thickness/width (nM) in TGFβ2 perfused tissue and reduction when supplemented with rhCLU. (*) indicates significance, with the median represented in a red line in the middle.

To establish the therapeutic potential of clusterin, we elucidated the role of clusterin in mitigating pathological IOP elevation induced by the profibrotic growth factor TGFβ2, a known inducer of ocular hypertension [30, 31]. The experimental paradigm involved baseline perfusion of DMEM in the human anterior segment perfusion model for 24 hours. This was followed by perfusion with rhTGFβ2 in DMEM for up to 150 hours (∼6 days) when the significant induction in IOP was achieved. Subsequently, rhCLU perfusion was carried out to identify if it mitigated the IOP elevation till 240 hours (∼10 days). Upon significant increase in IOP in response to TGFβ2 (p=0.02, n=4) by day 6 relative to the 24-hour baseline (**Figure 2B**), rhCLU at 0.5 μg/ml was added to the DMEM containing TGFβ2 till the conclusion of the experiment. The introduction of clusterin resulted in a gradual IOP reduction reaching statistical significance by the 216 hours (day 9) (p=0.008, n=4), with this significant decrease maintained through the 240th hour (day 10) (p=0.007, n=4) compared to the peak IOP level at 144 hours, coinciding with the start of clusterin supplementation, as depicted in **Figure 2B**. Thus, providing solid functional evidence for clusterin to counteract elevated IOP effectively. After terminating the experiment on day 10, tissues perfused with rhCLU, TGFβ2, and TGFβ2+rhCLU were fixed and processed for immunofluorescence and scanning electron microscopy (SEM).

### 3.3 **Clusterin induced antifibrogenic effects to lower TGFβ2-induced IOP elevation:**

Since the experiments were performed before and after rhCLU perfusion in the presence of TGFβ2, tissues perfused with rhCLU, TGFβ2, and the combination of TGFβ2+rhCLU were processed for EM and immunofluorescence analysis. The tissue perfused with rhCLU showed a decrease in the ECM proteins COL1 and fibronectin (FN) within the outflow pathway, including the TM and SC (**Figure 2C**). The top panel comprises COL1, and the bottom panel is FN (in green). Tissues perfused with TGFβ2 (middle panel) demonstrated an increase in COL1 and FN expression, which was mitigated by the perfusion of rhCLU (right panel).

For SEM imaging of the TM outflow pathway tissue was examined at a magnification of 10,000X. **Figure 2D** displays the representative images of the surface morphology indicating the structural fibrillar matrix of TM tissues treated with rhCLU (top), TGFβ2 (middle), and TGFβ2+rhCLU (bottom).

Treatment with TGFβ2 resulted in a noticeable thickening of this fibrillar matrix, marked by yellow arrows. In contrast, tissues treated with rhCLU, both alone and in combination with TGFβ2, exhibited a reduced thickening of the fibrillar matrix compared to those treated solely with TGFβ2. Using ImageJ for quantitative analysis, the thickness (width) of the structural fibrillar matrix was measured (**Figure 2E)**. In a blinded manner, the thickness measurements were obtained by discretizing the large image area into 10 imaginary subdivisions. One random fiber was chosen from each square by a non-experimenter for the measurement. This was repeated for 4 different tissues resulting in 40 measurements from each treatment condition which were coded. Post measurement, the codes were revealed, and the fibrillar matrix width was plotted as a violin plot with the median value highlighted in red. The fiber thickness was significantly greater in tissues treated with TGFβ2 alone (p=0.0001) compared to those treated with rhCLU. Conversely, the addition of rhCLU to TGFβ2-treated tissues significantly reduced the fibrillar matrix thickness (p=0.02) compared to TGFβ2 treatment alone. Put together, we found that secretory clusterin reduces IOP by increasing ECM remodeling in the TM.

### 3.4 **Secretory clusterin decreases actin contractility and cell adhesive interactions:**

Building on the observed impact of secretory clusterin on IOP and our earlier study showing that loss of clusterin function impacted IOP and actin cytoskeleton, we explored the underlying cellular and molecular basis for contributing to IOP lowering. In our pursuit to explore the mechanisms underlying cellular behavior, particularly the role of the actin-cytoskeleton, we performed a detailed study *in vitro* analysis on HTM cells. Serum-starved HTM cells were treated with 0.5 μg/ml of rhCLU for 2, 6, and 24 hours, and stained for actin using phalloidin (green) and the focal adhesion protein paxillin (red). To improve image clarity, both phalloidin and paxillin staining were digitally inverted using ImageJ. Compared to the untreated control, actin fibers were notably decreased by 6 and 24 hours (**Figure 3A**) with the paxillin staining intensity weakening at all the three time points studied (**Figure 3A**). An enhanced or zoomed view of the yellow-highlighted area in the projected image provided clear insight into the observed changes. Shown using red arrows in the middle panel indicate matured paxillin whose reduction under clusterin treatment was evident in a time-dependent manner. To derive quantitative data, we measured the F-actin and paxillin puncta staining intensities as well as counts were conducted from five distinct images across three biological replicates. Treatment of HTM cells with rhCLU led to a notable decrease in F-actin intensity relative to controls, with significant reductions observed after 6 hours (p=0.003) and 24 hours (p=0.001) of treatment (**Figure 3B**). Paxillin puncta intensity also decreased at 2 hours (p=0.04), 6 hours (p=0.009), and 24 hours (p=0.03) compared to controls, as shown in **Figure 3C**. Moreover, the decrease in the number of paxillin puncta, corresponding to reduced paxillin intensity, was noted across all time points with a significant reduction apparent after 24 hours of treatment (p=0.02) as depicted in **Figure 3D**. It was observed that clusterin treatment not only diminished F-actin organization but also reduced paxillin alignment along actin fibers in comparison to the control.

**Figure 3:**
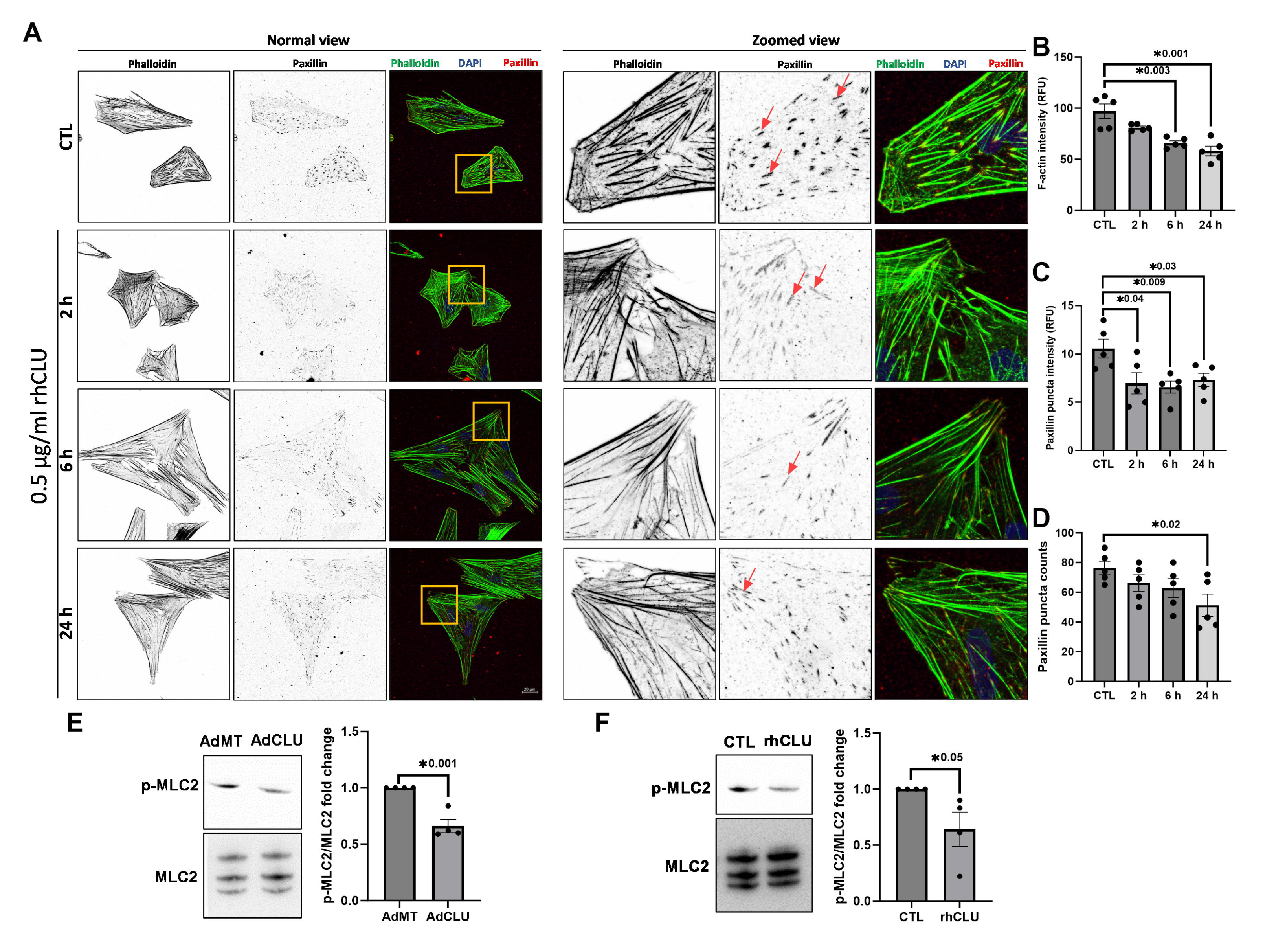
Clusterin decreases actin contractility and cell adhesion interactions. A) HTM cells treated with 0.5 μM rhCLU at 2 h, 6, and 24 h showed a reduction in F-actin fibers stained by phalloidin in green and distribution of focal adhesion protein paxillin in red. For better visualization, individual layouts of phalloidin and paxillin are inverted using Image J. A Zoomed view of selected portions is provided on the right, with red arrows in the magnified middle panel indicating matured paxillin. The nucleus is stained with DAPI in blue. The scale bar is 20 μm. To the far right is a graphical representation of changes in F-actin intensity (B), paxillin puncta intensity (C), and paxillin puncta counts (D), which are provided across treatment conditions. E) Immunoblot (IB) analysis of the ratio of p- MLC2/MLC2 in PTM cells treated with AdCLU compared to AdMT. The graphical representation shows a reduction in p-MLC2 in AdCLU compared to AdMT. F) IB analysis of the ratio of p-MLC2/MLC2 in PTM cells treated with rhCLU compared to control. The graphical representation shows a reduction in p- MLC2 in rhCLU compared to control. Error bars represent the SEM, and (*) indicates significance, with a sample size of n=3.

Paxillin acts as a cytoskeleton scaffolding protein at focal adhesion sites, where it interacts with integrins, actin filaments, regulators of Rho GTPase, and other focal adhesion components such as vinculin, talin, and focal adhesion kinase. Changes in paxillin expression or phosphorylation state can alter the dynamics of focal adhesions, thereby affecting cell adhesion strength, and cellular contraction [9, 32].

A defining feature of TM cells is their contractile nature governed by the actomyosin changes alongside focal adhesion proteins such as paxillin. Notably, an increase in IOP has been linked to enhanced TM cell contractility via phosphorylation of MLC2 [9]. As we observed a reduction in actin fibers, paxillin counts, and their intensity following clusterin treatment, we investigated the phosphorylation status of MLC2 (p- MLC2) under both AdCLU and rhCLU in PTM cells. **Figures 3E** and **3F** display the ratio of p-MLC2 to MLC2 under conditions of clusterin induction via AdCLU and rhCLU, respectively. Both approaches resulted in a significant reduction in the p-MLC2/MLC2 ratio - AdCLU (p=0.001, n=4) (**Figure 3E)** and rhCLU (p=0.05, n=4) (**Figure 3F**) compared to their control. Thus, defining the onset of cellular relaxation in TM due to an increase in clusterin availability and providing a mechanistic basis for the IOP reduction seen with clusterin treatment.

### 3.5 **Differential protein expression in HTM due to constitutive clusterin expression reveals altered protein quality control and cellular biomechanics.**

#### Proteins upregulated by constitutive clusterin expression

The alterations observed in the actin cytoskeleton, focal adhesion proteins, and cell contractility following clusterin treatment led us to utilize comprehensive and unbiased global proteomics to delineate the molecular pathways using HTM cells treated with AdCLU or AdMT. From a TMT-based LC/MS-MS proteomics approach, 5,647 proteins were identified to be changed and 5162 protein changes were quantified. Based on FDR ≤5%, statistical significance (p ≤ 0.050), and mean ± 2σ of log2 ≥ 0.1 of confidence fold change limits, 214 proteins were significantly upregulated and 150 were significantly downregulated in TM under AdCLU treatment compared to AdMT. **Figure 4** showcases the heatmap for the top 60 proteins that were either upregulated or downregulated, using a color gradient from blue to red with blue indicating lower expression and red indicating higher expression levels. The list of top 10 upregulated proteins is provided in **Table 3**.

**Figure 4:**
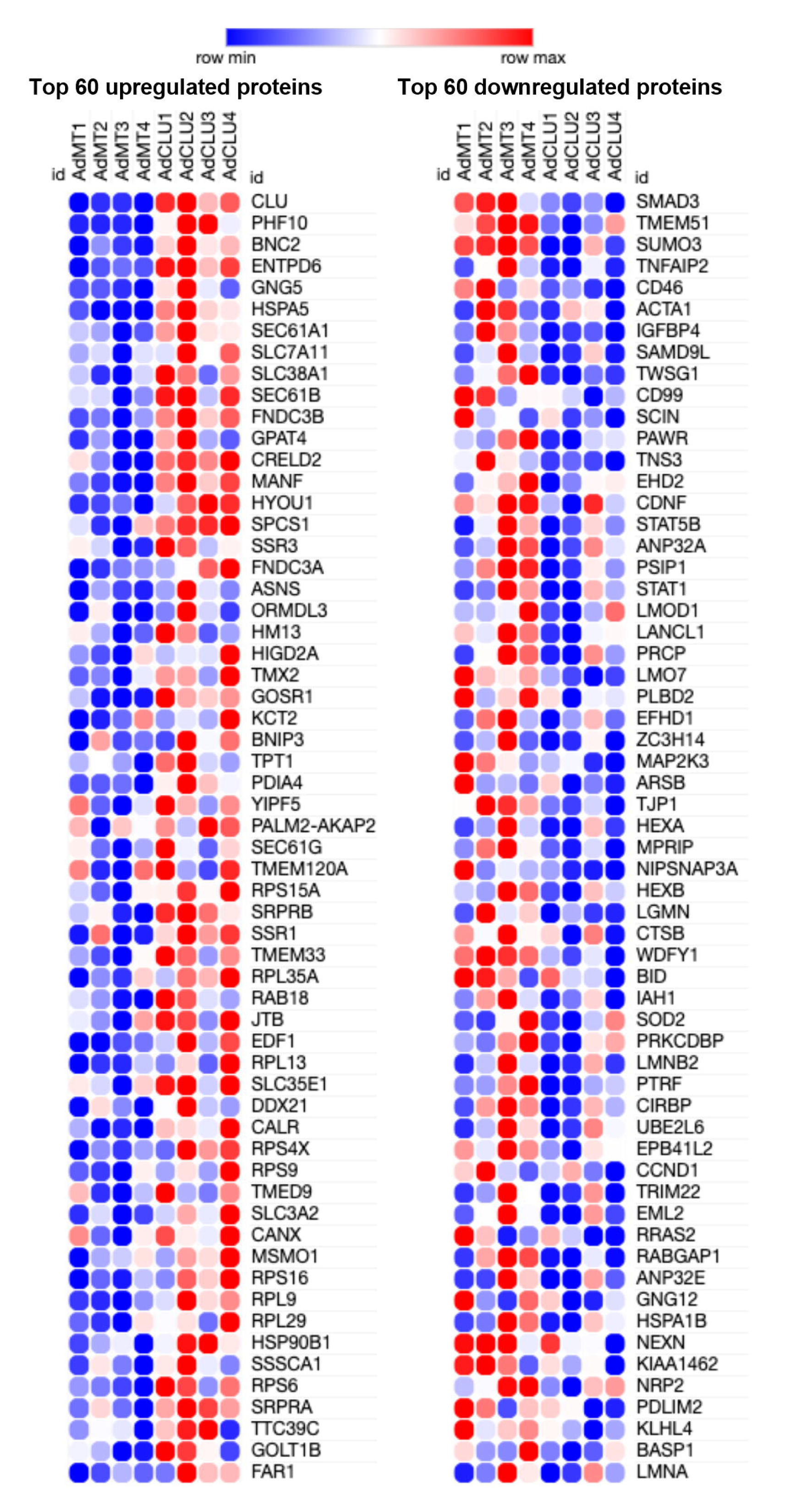
Heat map of differentially expressed proteins upon clusterin induction. Heatmap for the top 60 proteins that were either upregulated or downregulated by AdCLU, using a color gradient from blue to red to denote expression levels: with blue indicating lower expression (row min) and red indicating higher expression (row max) levels.

**Table 3:**
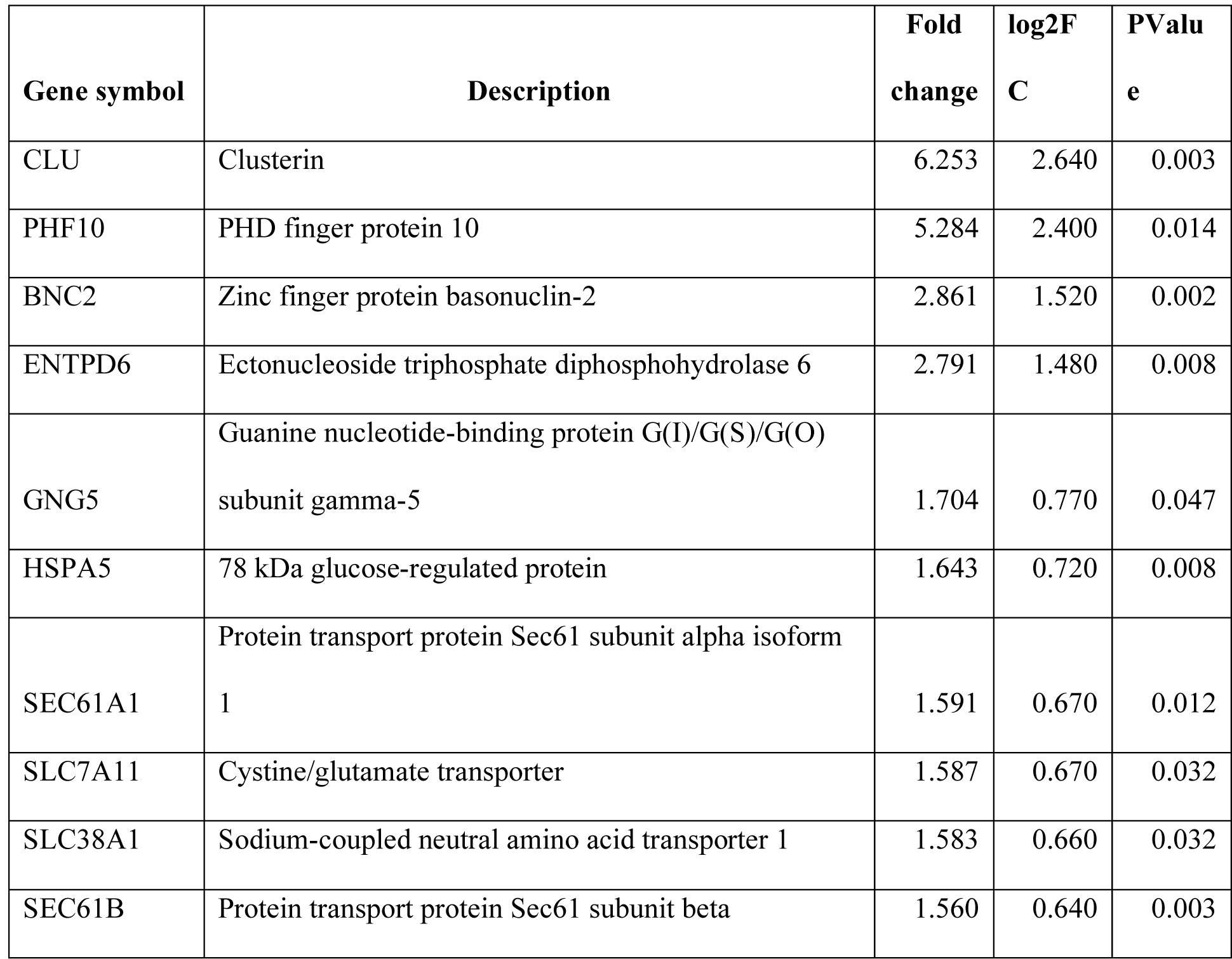
List of top 10 upregulated proteins.

Under AdCLU, the most significantly upregulated protein was clusterin (2.64-log2FC; 6.25-FC) validating the experimental paradigm. This was followed by – **a)** the PHD finger protein 10 (PHF10) (2.4-log2FC; 5.28-FC) involved in transcription activity regulation by chromatin remodeling [33]. PHF10 is part of polybromo-associated BAF (PBAF) complexes that have been reported to be involved in maintaining genomic stability and have a role in regulating chromatin architecture [34]. **b)** Zinc finger protein basonuclin-2 (BNC2) (1.52-log2FC; 2.86-FC) that controls the expression of collagens, matrix metalloproteases (MMPs), and other matrisomal components in cancer cells [35]. **c)** membrane protein ectonucleoside triphosphate diphosphohydrolase 6 (ENTPD6) (1.48-log2FC; 2.79-FC), which is an enzyme involved in catalyzing the hydrolysis of nucleoside di- and triphosphate [36]. **d)** guanine nucleotide-binding protein G(I)/G(S)/G(O) subunit gamma-5 (GNG5), a G protein (0.77-log2FC; 1.70- FC). **e)** proteins involved in amino acid and peptide transport within cellular components - the cystine/glutamate transporter (SLC7A11) (0.67-log2FC; 1.59-FC) and the sodium-coupled neutral amino acid transporter (SLC38A7) (0.66-log2FC; 1.58-FC) – aid in the regulation of amino acid levels within cells, impacting cellular metabolism and the antioxidant response [37, 38]. The Sec61 subunits - protein transport protein Sec61 subunit alpha isoform 1 (SEC61A1) (0.67-log2FC; 1.58-FC), protein transport protein Sec61 subunit beta (SEC61B) (0.64-log2FC; 1.56-FC), protein transport protein Sec61 subunit gamma (SEC61G) (0.48-log2FC; 1.39-FC). They are part of the machinery that transports proteins into the ER, an essential step in the synthesis and sorting of membrane and secretory proteins. **f)** Proteins critical for cellular quality control, which are involved in promoting protein folding, trafficking, and preventing the aggregation of unfolded proteins. This group includes heat shock proteins like the 78 kDa glucose-regulated protein (HSPA5) (0.72-log2FC; 1.64-FC), endoplasmin (0.39-log2FC; 1.30-FC), stress- 70 protein, mitochondrial (HSPA9) (0.33-log2FC; 1.26-FC), DnaJ homolog subfamily B member 12 (DNAJB12) (0.21-log2FC; 1.15-FC). Additionally, other chaperone proteins like hypoxia up-regulated protein 1 (HYOU1) (0.56-log2FC; 1.47-FC), calreticulin (CALR) (0.42-log2FC; 1.34-FC) showed increased levels. Enzymes involved in ubiquitination processes, including E3 ubiquitin-protein ligase RNF185 (RNF185) (0.31-log2FC; 1.24-FC), E3 ubiquitin-protein ligase CHIP (STUB1) (0.27-log2FC; 1.21-FC), ubiquitin-conjugating enzyme E2 R2 (UBE2R2) (0.15-log2FC; 1.11-FC), ubiquitin-conjugating enzyme E2 G2 (UBE2G2) (0.11-log2FC; 1.08-FC) were also significantly upregulated. We propose the increase in could be to increase the production of clusterin and secrete it. **g)** cytoskeleton-related proteins, including thymosin beta-10 (0.25-log2FC; 1.19-FC), known for its role in actin sequestration [39], and drebrin (0.22-log2FC; 1.17-FC) which modulates actomyosin interaction [40] and Protein furry homolog-like (0.12-log2FC; 1.089-FC) that regulates actin-cytoskeleton. **h)** some of the ECM and ECM-associated proteins such as fibronectin type III domain-containing protein 3A (FNDC3A) (0.53-log2FC; 1.45-FC) and 3B (FNDC3B) (0.64-log2FC; 1.56-FC), which are involved in cell adhesion and ECM assembly [41], along with collagen alpha-1(V) chain (COL5A1) (0.34-log2FC; 1.26-FC). **i**) Proteins involved in collagen biosynthesis prolyl 4-hydroxylase subunit alpha-2 (P4HA2) (0.25-log2FC; 1.19-FC) and Peptidyl-prolyl cis-trans isomerase FKBP14 (FKBP14) (0.24-log2FC; 1.18- FC). **j**) Additionally, integrin alpha-5 (ITGA5) (0.26-log2FC; 1.20-FC), linking ECM with cytoskeleton, and mucin-15 (MUC15) (0.29-log2FC; 1.22-FC), a cell adhesion protein.

#### Pathway enrichment analysis for proteins upregulated by constitutive clusterin expression

Pathway enrichment analysis using ShinyGO [42] for upregulated proteins in TM induced by clusterin based on molecular function and cellular component are given in **Table 4** and **Supplementary Table 2** respectively. The enriched pathways predominantly include proteins associated with structural molecule activity, such as those involved in ribosomal assembly and function, suggesting an upregulation of protein synthesis. Intriguingly, the analysis also highlighted proteins with small molecule binding properties, which are often integral to the stress response and quality control mechanisms, pointing to a cellular adaptation aimed at preserving homeostasis.

**Table 4:**
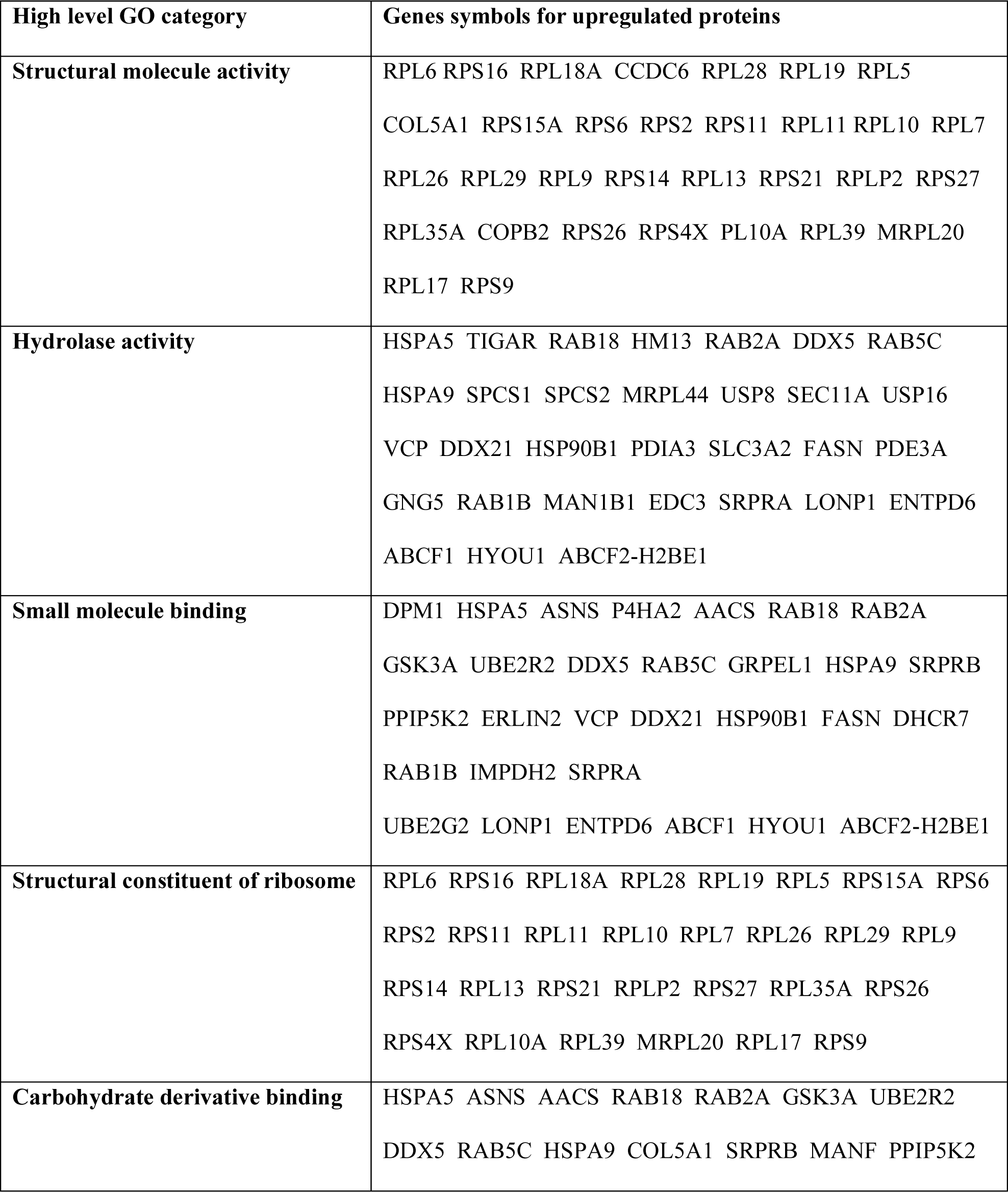
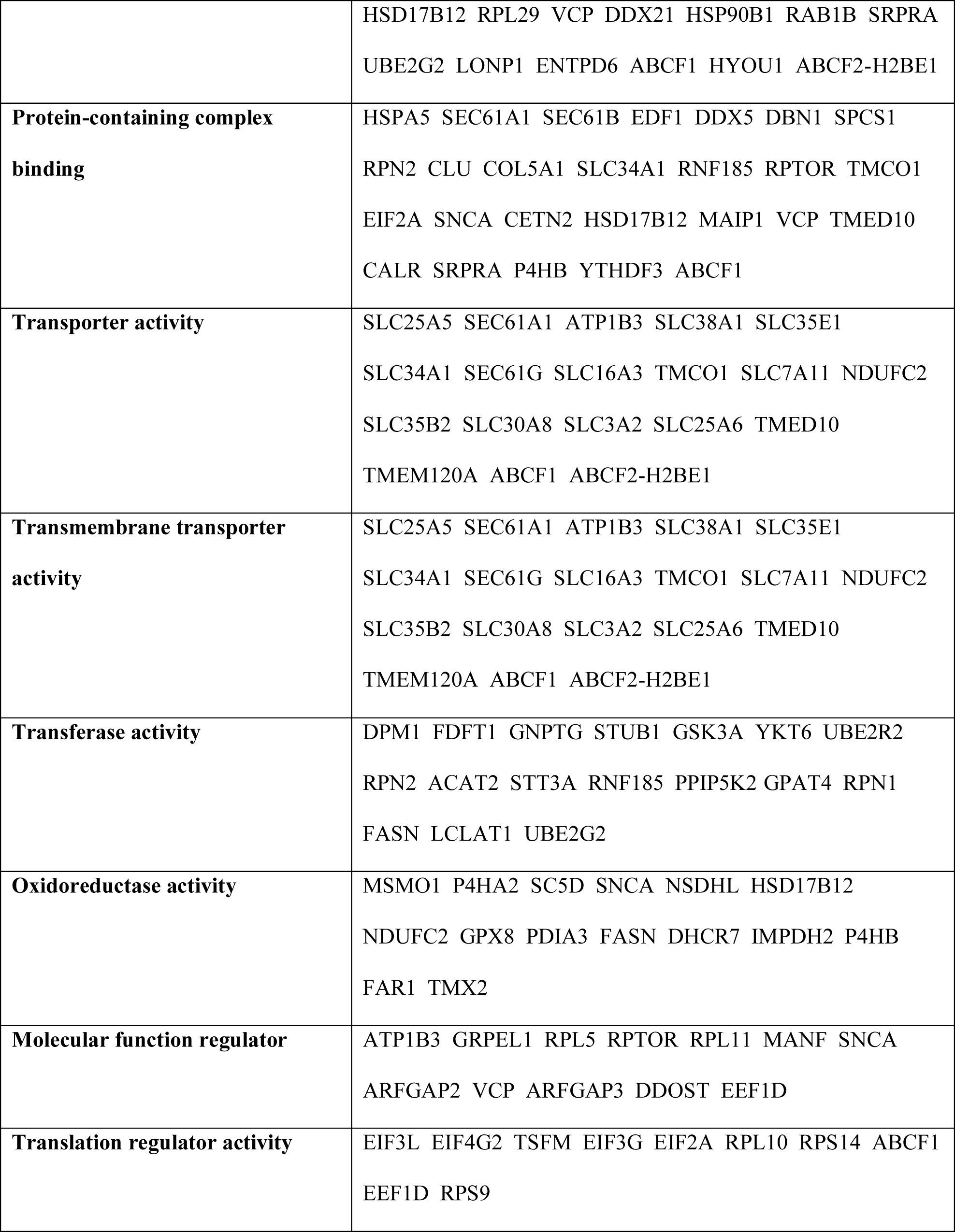
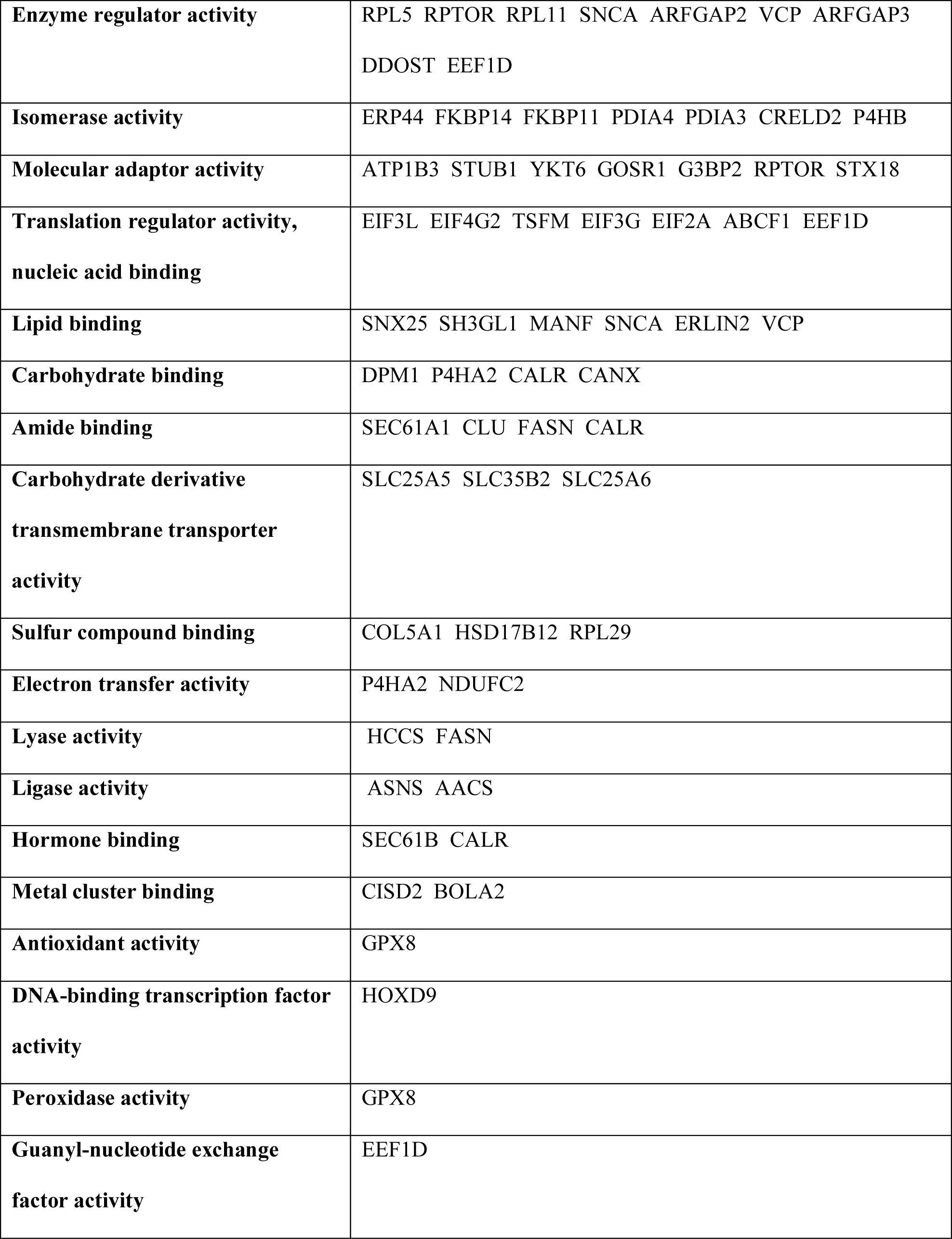
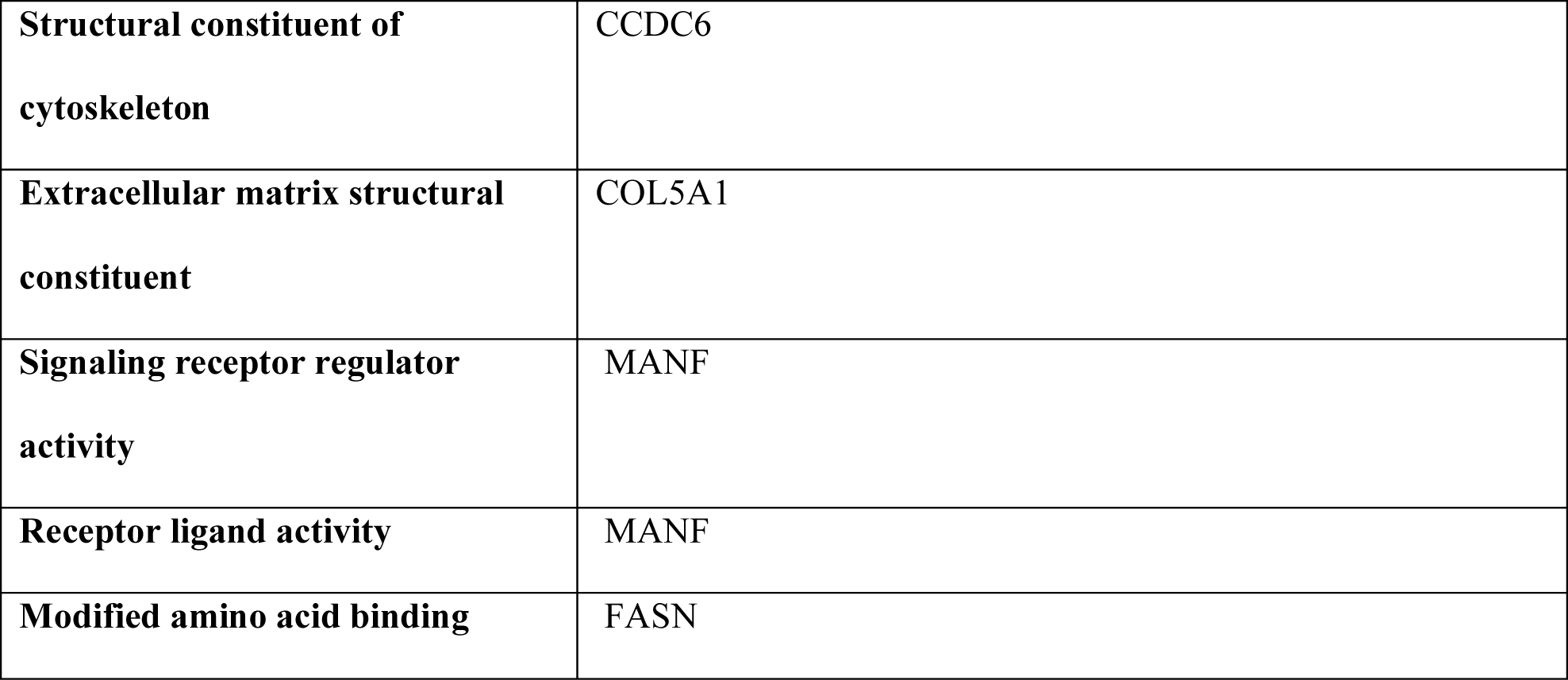
Pathway enrichment analysis for upregulated cellular proteins using ShinyGO based on molecular function.

#### Proteins downregulated by constitutive clusterin expression

Among the most downregulated proteins, the top 10 are provided in **Table 5**. Of great physiological significance are the following – **a**) Mothers against decapentaplegic homolog 3 (SMAD3) (-0.9-log2FC; 0.54-FC), which is a part of TGFβ2 canonical signaling [43] and TGFβ2 induced IOP elevation is dependent on SMAD3 [14]. **b**) Transmembrane protein 51 (TMEM51) (-0.72-log2FC; 0.61-FC) potentially involved in interacting with actin regulators like Cdc42 and FERM and PDZ domain-containing protein1. **c**) small ubiquitin-related modifier 3 (SUMO3) (-0.70-log2FC; 0.61-FC) potentially function to antagonize ubiquitin, small ubiquitin-related modifier 1 (SUMO1) (-0.26-log2FC; 0.84-FC) involved in ubiquitination, ubiquitin associated proteins like Ubiquitin/ISG15-conjugating enzyme E2 L6 (UBE2L6) (-0.32-log2FC; 0.80-FC), and E3 ubiquitin-protein ligase TRIM22 (TRIM22) (-0.32-log2FC; 0.80-FC), indicating a potential regulatory role of clusterin in cellular ubiquitination and SUMOylation processes, impacting various protein functions, and cellular stress responses. **d**) Of significance, clusterin downregulated a spectrum of proteins that play key roles in cytoskeletal dynamics, cell adhesive and cell-matrix interactions, and cell junctions, emphasizing the impact of clusterin on cellular structure and function. Clusterin significantly decreased alpha skeletal muscle actin (ACTA1) (-0.51-log2FC; 0.70-FC) essential for cytoskeletal structure; adseverin (SCIN) (-0.45-log2FC; 0.73-FC) that modulates actin filament dynamics; leiomodin-1 (LMOD1) (-0.40-log2FC; 0.76-FC) involved in actin nucleation; gelsolin (GSN) (-0.20-log2FC; 0.87-FC) known for severing and capping actin filaments; and Wiskott-Aldrich syndrome protein family member 2 (WASF2) (-0.14-log2FC; 0.91- FC), associated with actin assembly; palladin (PALLD) (-0.19-log2FC; 0.88-FC) that is key to the organization of actin filaments; alpha-actinin-4 (ACTN4) (-0.17-log2FC; 0.89-FC) that is important for F-actin cross-linking and contributing to cellular strength and integrity; filamin-binding LIM protein 1 (FBLIM1) (-0.14-log2FC; 0.91-FC), which plays a role in linking the actin cytoskeleton to the cell membrane; and plectin (PLEC) (-0.20-log2FC; 0.87-FC) that is critical in connecting various cytoskeletal networks and providing mechanical resilience to cells. **e**) Focal adhesion proteins tensin 3 (-0.44-log2FC; 0.74-FC) and tensin 4 (-0.22-log2FC; 0.86-FC), which link the ECM to the actin cytoskeleton through integrins and cell surface receptors influencing cytoskeletal organization. Although tensin 1 downregulation (-0.23-log2FC; 0.86-FC) did not fall under the criteria of significance, it exhibited a notable reduction. Additionally, the tight junction protein ZO-1 (TJP1) (-0.370-log2FC; 0.78-FC), myosin phosphatase Rho-interacting protein (MPRIP) (-0.360-log2FC; 0.78-FC), myosin regulatory light chain 12B (MYL12B) (-0.13-log2FC; 0.91-FC), and caldesmon (CALD1) (-0.25-log2FC; 0.84-FC), a key regulator of actin-myosin interactions indicating the potential negative regulation of cell-cell adhesion and cellular contractility due to increasing clusterin levels. **f**) Moreover, the downregulation of proteins involved in signaling pathways such as serine/threonine-protein kinase N2 (PKN2) (-0.12-log2FC; 0.92- FC) and Rho-related GTP-binding protein RhoQ (RHOQ) (-0.12-log2FC; 0.92-FC) highlights the broad scope of clusterin, extending beyond structural components to encompass key elements in the cytoskeletal signaling network. Additionally, the reduction in echinoderm microtubule-associated protein-like 2 (EML2) (-0.31-log2FC; 0.81-FC) highlighted changes in tubulin cytoskeleton stability. **g**) signaling molecules like signal transducer and activator of transcription (STAT) family members, with STAT1 (- 0.40-log2FC; 0.76-FC), STAT3 (-0.23-log2FC; 0.85-FC), and STAT5B (-0.43-log2FC; 0.74-FC). STAT2 also exhibited a decrease (-0.34-log2FC; 0.79-FC), although it did not meet the significant threshold. **h**) G proteins including guanine nucleotide-binding protein G(I)/G(S)/G(O) subunit gamma-12 (GNG12) (- 0.29-log2FC; 0.82-FC) and guanine nucleotide-binding protein G(I)/G(S)/G(T) subunit beta-1 (GNB1) (- 0.21-log2FC; 0.87-FC); and ras-related protein R-Ras2 (RRAS2) (-0.3-log2FC; 0.81-FC), a member of the Ras GTPase.

**Table 5:** List of top 10 downregulated proteins.

| **Gene**<br>**symbol** | **Description** | **Fold**<br>**change** | **log2FC** | **PValue** |
| --- | --- | --- | --- | --- |
| SMAD3 | Mothers against decapentaplegic homolog 3 | 0.536 | -0.900 | 0.005 |
| TMEM51 | Transmembrane protein 51 | 0.608 | -0.720 | 0.029 |
| SUMO3 | Small ubiquitin-related modifier 3 | 0.614 | -0.700 | 0.011 |
| TNFAIP2 | Tumor necrosis factor alpha-induced protein 2 | 0.694 | -0.530 | 0.033 |
| CD46 | Membrane cofactor protein | 0.696 | -0.520 | 0.024 |
| ACTA1 | Actin, alpha skeletal muscle | 0.700 | -0.510 | 0.046 |
| IGFBP4 | Insulin-like growth factor-binding protein 4 | 0.701 | -0.510 | 0.041 |
| TWSG1 | Twisted gastrulation protein homolog 1 | 0.713 | -0.490 | 0.039 |
| SAMD9L | Sterile alpha motif domain-containing protein 9-like | 0.714 | -0.490 | 0.011 |
| CD99 | CD99 antigen | 0.717 | -0.480 | 0.021 |

#### Pathway enrichment analysis for proteins downregulated by constitutive clusterin expression

Pathway enrichment analysis using ShinyGO [42] for downregulated proteins in TM induced by clusterin based on molecular function and cellular component are given in **Table 6** and **Supplementary Table 3** respectively. The analysis revealed that the molecular functions of the majority of the downregulated proteins in TM after inducing clusterin expression were linked to hydrolase activity, GPCR signaling, structural constituents of the cytoskeleton, and MAPK kinase activity. These findings define the significant impact of clusterin on cellular structure and dynamics.

**Table 6:**
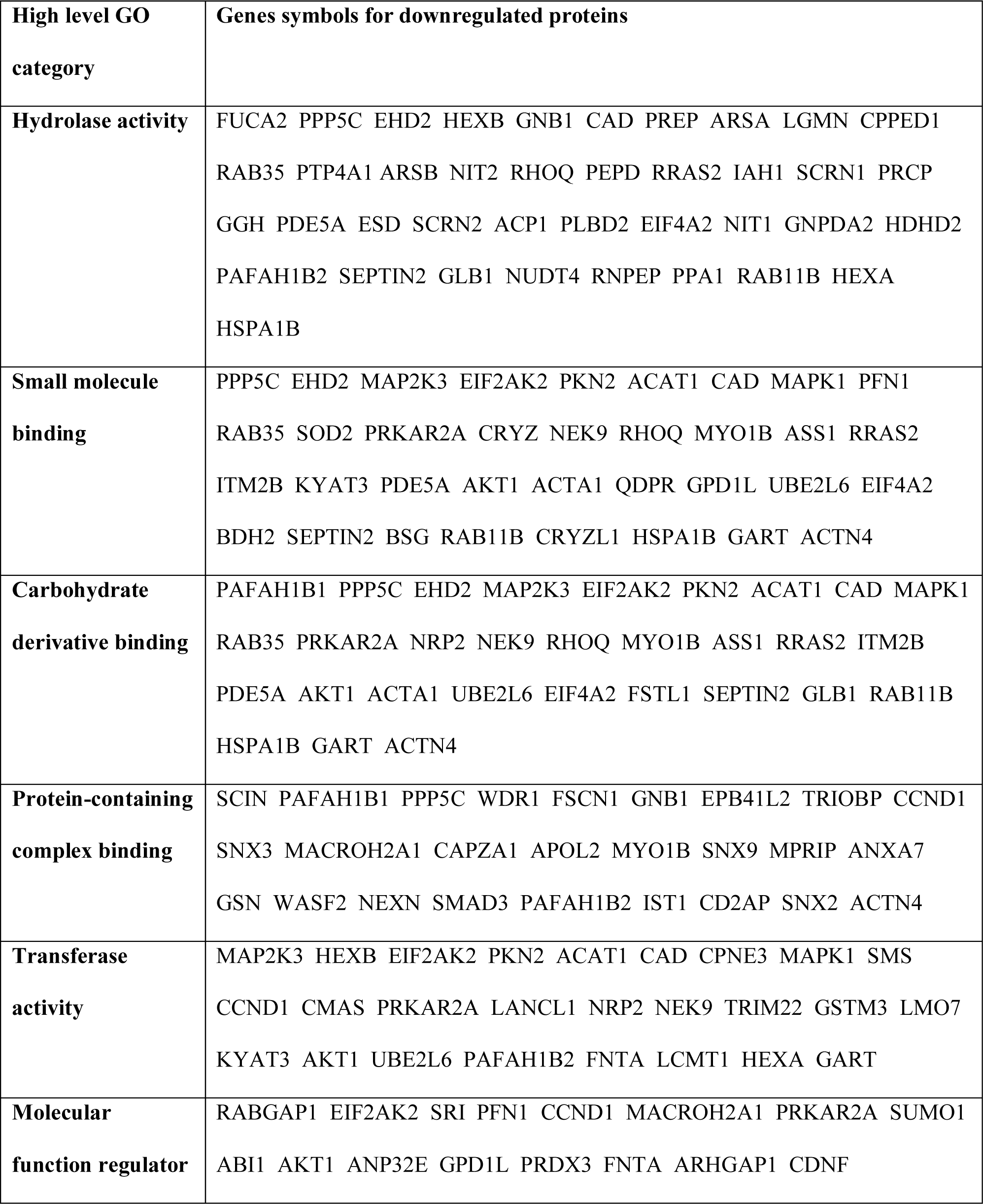
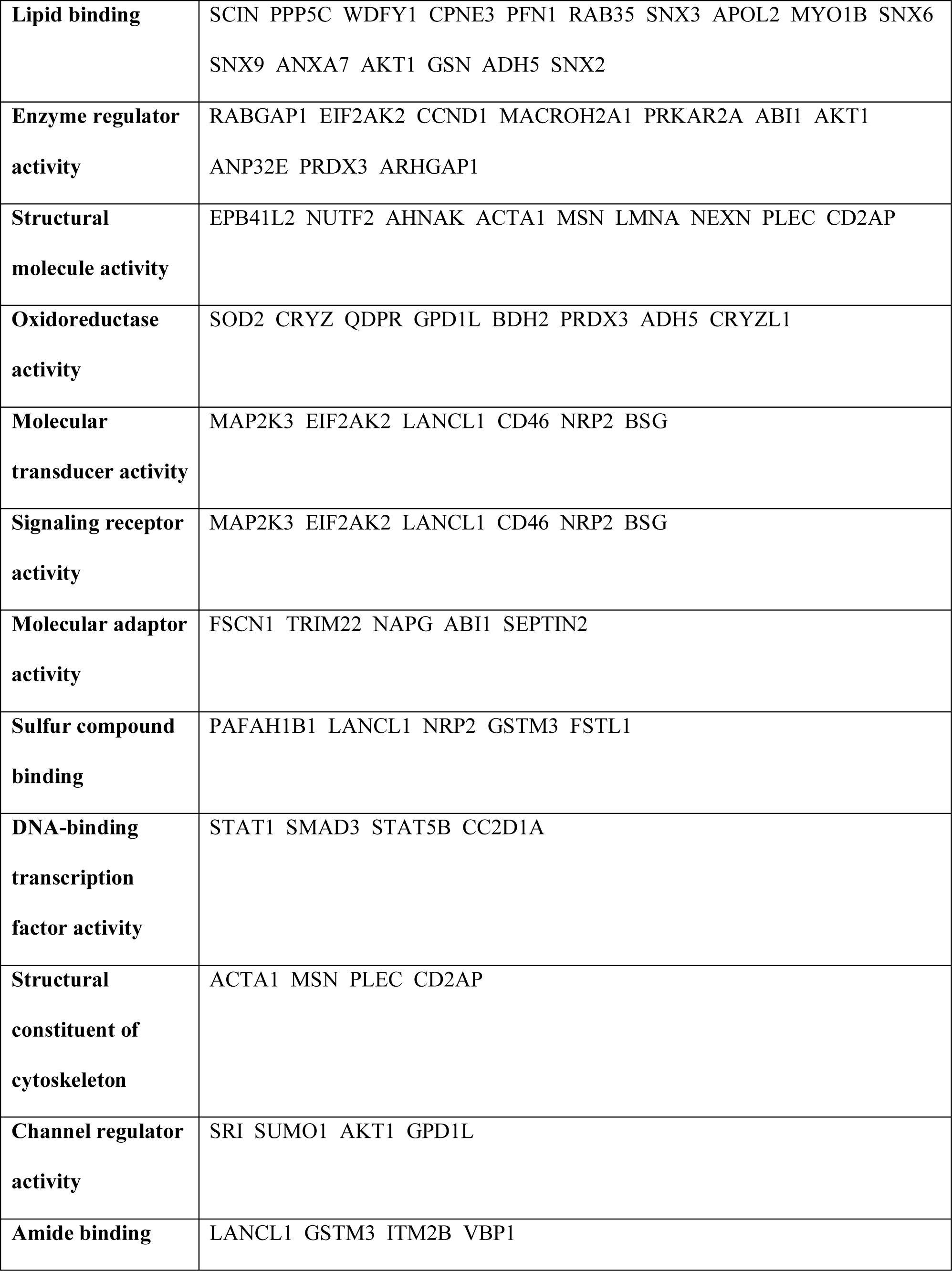
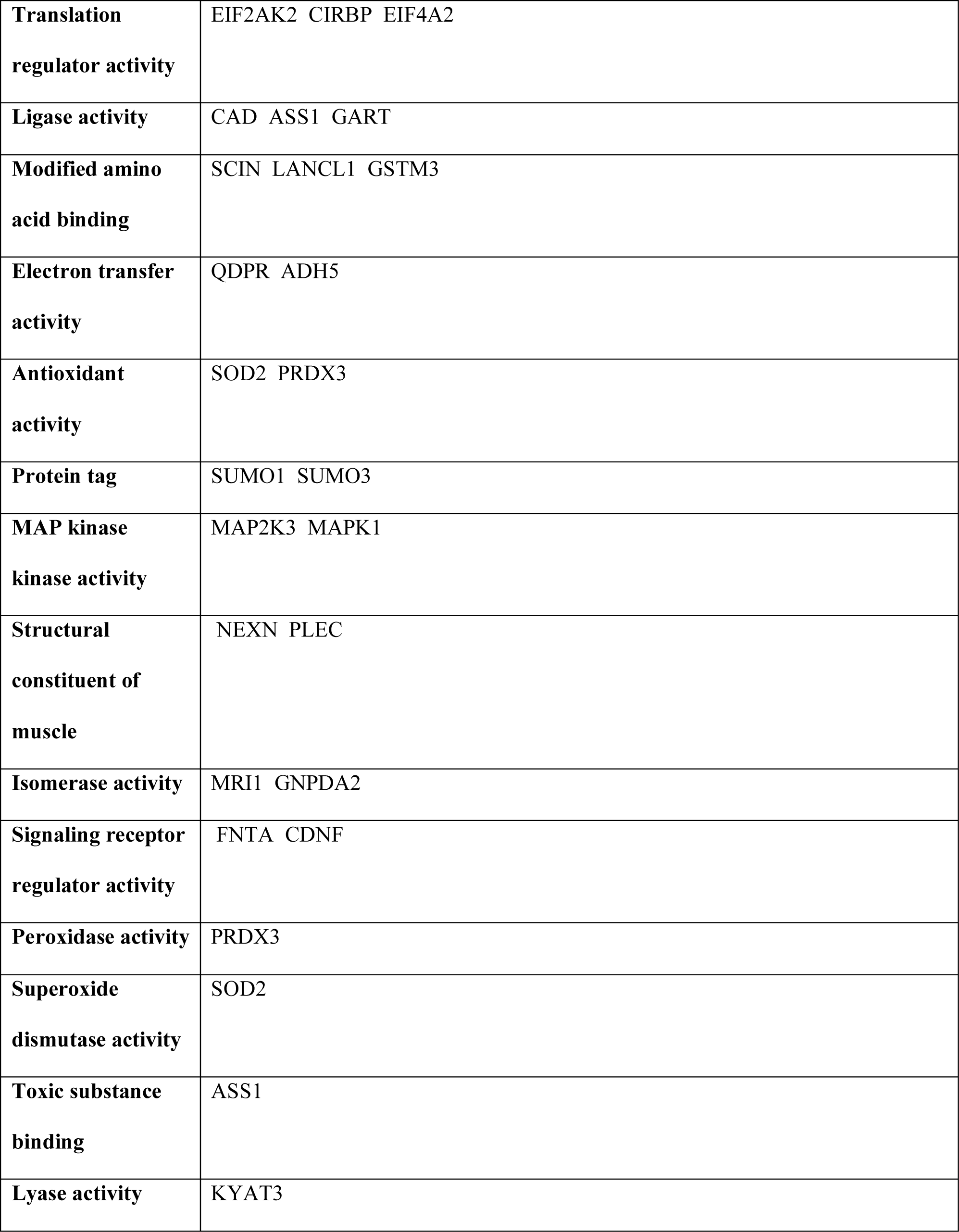
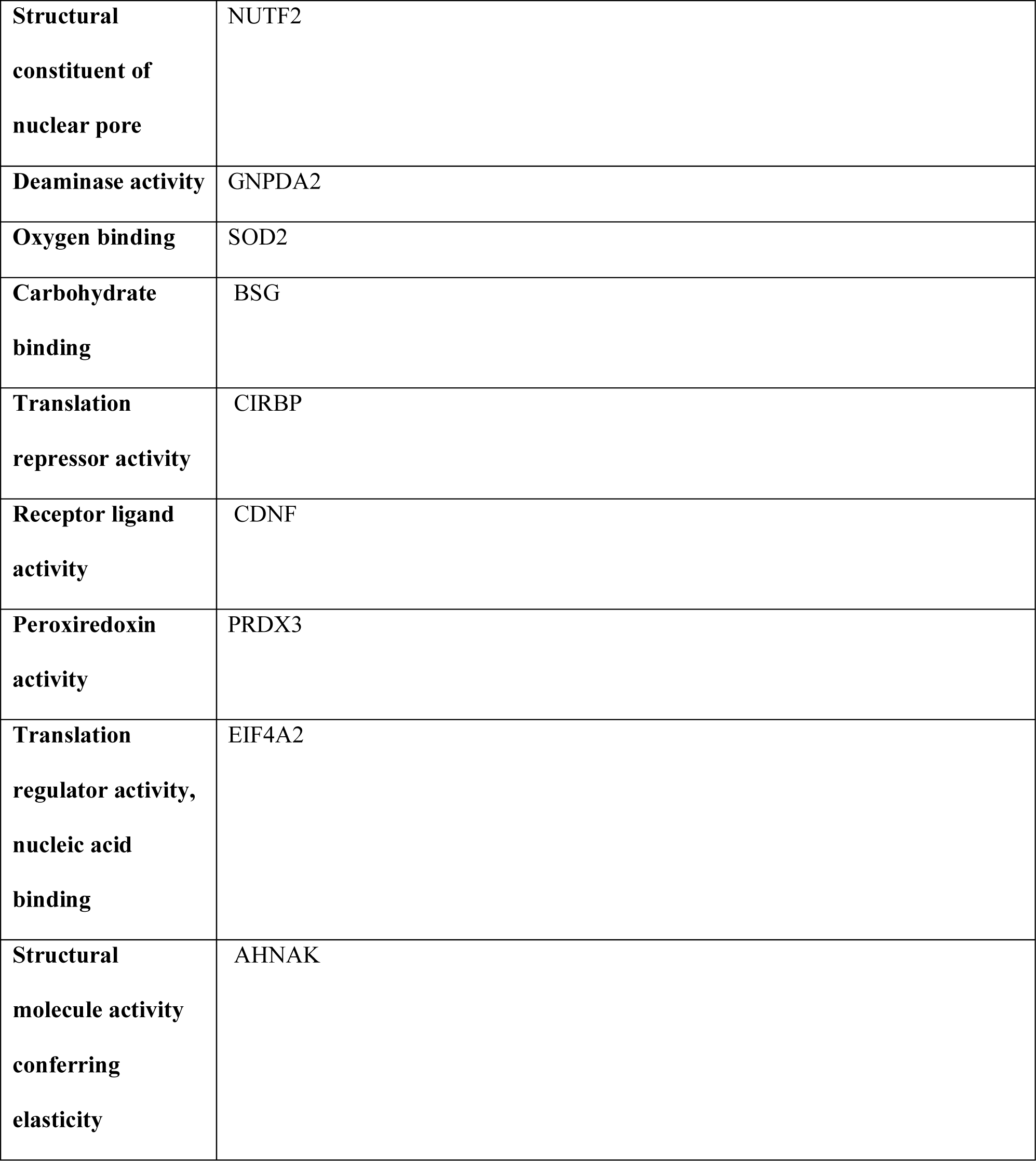
Pathway enrichment analysis for downregulated cellular proteins using ShinyGO based on molecular function.

### 3.6 **Proteomic analysis of the ECM enriched fraction reveals the association of clusterin to ECM.**

Since we observed significant changes in the actin cytoskeleton, cell-adhesive interactions, and cell-matrix interactions upon AdCLU treatment, we narrowed down our focus to ECM and ECM-related changes. Using TMT-based LC/MS-MS proteomics on the ECM-enriched samples after AdCLU treatment identified 2,616 proteins and 2,238 were quantified. Further, based on FDR ≤5% and statistical significance (p ≤ 0.050), 45 proteins were significantly upregulated and 20 were significantly downregulated. Based on mean ± 2σ of log2 ≥ 0.1 of confidence fold change limits, 24 proteins were upregulated, and 10 proteins were downregulated. Further, using the criteria mean ± 2σ of log2 < 0.3–0.1, 21 proteins were upregulated, and 10 proteins were downregulated in TM under AdCLU treatment compared to AdMT.

The top 10 upregulated proteins are provided in **Table 7**. In the course of characterizing our ECM-enriched samples, we observed not only the expected ECM components but also a moderate presence of proteins typically associated with nuclear and cytoplasmic compartments. The presence of these proteins at moderate levels suggests they might be co-enriched, possibly due to their interaction with ECM components or due to cellular remnants that were not completely removed in the enrichment process.

**Table 7:** List of top 10 upregulated proteins in ECM enriched fraction.

| **Gene**<br>**symbol** | **Description** | **Fold**<br>**change** | **log2FC** | **PValue** |
| --- | --- | --- | --- | --- |
| CLU | Clusterin | 5.585 | 2.480 | 0.050 |
| HR | Lysine-specific demethylase hairless | 3.072 | 1.620 | 0.028 |
| ACOT8 | Acyl-coenzyme A thioesterase 8 | 1.869 | 0.900 | 0.019 |
| CDV3 | Protein CDV3 homolog1 | 1.594 | 0.670 | 0.044 |
| SRSF7 | Serine/arginine-rich splicing factor 7 | 1.553 | 0.640 | 0.038 |
| LDHB | L-lactate dehydrogenase B chain | 1.498 | 0.580 | 0.000 |
| STRIP2 | Striatin-interacting protein 2 | 1.492 | 0.580 | 0.027 |
| ARMCX3 | Armadillo repeat-containing X-linked protein 3 | 1.433 | 0.520 | 0.042 |
| NDUFA7 | NADH dehydrogenase [ubiquinone] 1 alpha subcomplex<br>subunit 7 | 1.427 | 0.510 | 0.035 |
| MANF | Mesencephalic astrocyte-derived neurotrophic factor | 1.394 | 0.480 | 0.042 |

Moreover, the application of DNase at a concentration lower than ideal in our protocol might have also contributed to the residual presence of these proteins.

Clusterin emerged as the most significantly upregulated protein (2.48-log2FC; 5.59-FC), transcriptional regulator lysine-specific demethylase hairless (HR) (1.62-log2FC; 3.07-FC), acyl-coenzyme A thioesterase 8 (ACOT8) (0.9-log2FC; 1.87-FC) involved in lipid metabolism, collagen alpha-1(III) chain (COL3A1) (0.43-log2FC; 1.34-FC), exostosin-1 (EXT1) (0.23-log2FC; 1.17-FC) contributing in heparan-sulfate synthesis [44], and procollagen-lysine,2-oxoglutarate 5-dioxygenase 2 (PLOD2) (0.35-log2FC; 1.28-FC) that stabilizes collagen cross-link [45]. Significant increases were also observed in proteins linked to cytoskeletal integrity and architecture. Striatin-interacting protein 2 (STRIP2) (0.58-log2FC; 1.49-FC) is involved in regulating cell morphology and cytoskeletal organization [46], protein 4.1 (EPB41) (0.23-log2FC; 1.18-FC) regulating spectrin-actin interaction contributing to cellular mechanical stability [47], and alpha-taxilin (TXLNA) (0.28-log2FC; 1.22-FC), involved in microtubule organization [48].

Among the downregulated proteins (**Table 8**) were - tRNA-splicing ligase RtcB homolog (RTCB) (-0.62- log2FC; 0.65-FC) involved in RNA maturation, cytoskeletal intermediate filament protein keratin, type I cytoskeletal 18 (KRT18) (-0.34-log2FC; 0.79-FC), CD2-associated protein (CD2AP) (-0.14-log2FC; 1.1- FC), an adaptor protein between the actin-cytoskeleton and membrane proteins [49].

**Table 8:** List of top 10 downregulated proteins in ECM enriched fraction.

| **Gene**<br>**symbol** | **Description** | **Fold**<br>**change** | **log2FC** | **PValue** |
| --- | --- | --- | --- | --- |
| RTCB | tRNA-splicing ligase RtcB homolog | 0.651 | -0.620 | 0.047 |
| ARMH2 | Armadillo-like helical domain-containing protein 2 | 0.660 | -0.600 | 0.020 |
| SF1 | Splicing factor 1 | 0.667 | -0.580 | 0.042 |
| RABGAP1 | Rab GTPase-activating protein 1 | 0.693 | -0.530 | 0.019 |
| GNG12 | Guanine nucleotide-binding protein G(I)/G(S)/G(O) subunit<br>gamma-12 | 0.695 | -0.520 | 0.049 |
| RBMX | RNA-binding motif protein, X chromosome | 0.708 | -0.500 | 0.019 |
| H2AFX | Histone H2AX | 0.750 | -0.420 | 0.035 |
| CD55 | Complement decay-accelerating factor | 0.786 | -0.350 | 0.049 |
| KRT18 | Keratin, type I cytoskeletal 18 | 0.792 | -0.340 | 0.012 |
| NSDHL | Sterol-4-alpha-carboxylate 3-dehydrogenase,<br>decarboxylating | 0.795 | -0.330 | 0.041 |

The guanine nucleotide-binding protein G(q) subunit alpha (GNAQ) (-0.14-log2FC; 0.91-FC) was significantly decreased. In line with our observations from the whole cell lysate proteomics data, we noted a significant reduction in the levels of the G protein GNG12 (-0.52-log2FC; 0.70-FC) and the Ras-related protein RRAS2 (-0.14-log2FC; 0.91-FC). These clusterin-induced changes in GNG12, GNAQ, and RRAS2 suggest alterations in G protein signaling and Ras-related pathways, potentially impacting cellular communication and response mechanisms.

String network analysis (**Figure 5)** maps the interactions between proteins associated with the actin cytoskeleton, G protein signaling, and ubiquitination pathways, encompassing both upregulated and downregulated proteins following clusterin induction. Notably, the actin-associated proteins that were significantly downregulated by clusterin exhibited a high degree of interconnectivity within the analysis. Similarly, the network shows a robust interconnection among all proteins associated with G protein signaling that were downregulated.

**Figure 5:**
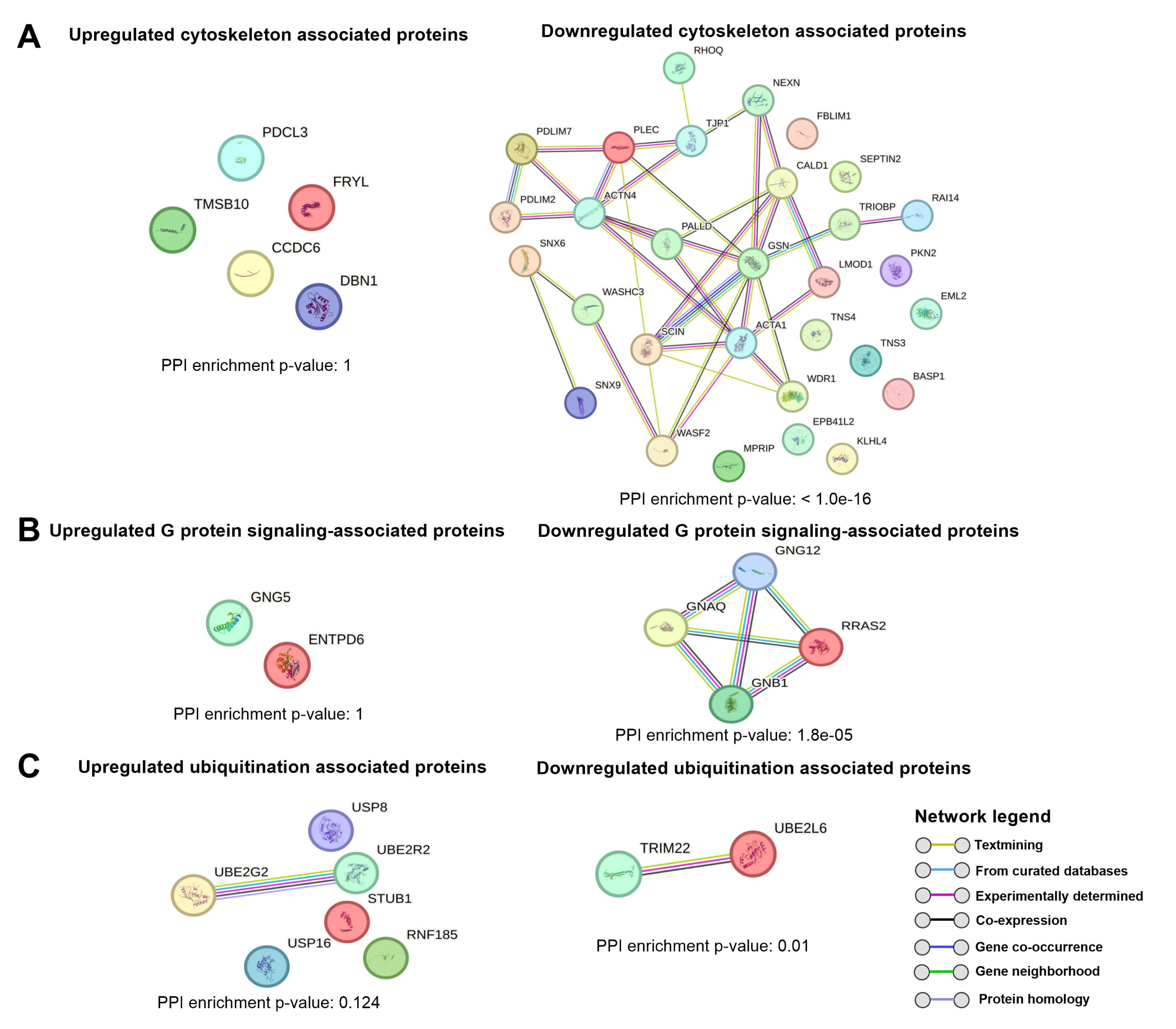
String network analysis elucidates the alteration in protein interactions related to actin-cytoskeleton, G proteins, and ubiquitination. STRING network analysis of upregulated and downregulated (A) actin-cytoskeleton, (B) G proteins, and (C) ubiquitination-associated proteins. Colored lines between the nodes indicate their basis for interconnection. Overall PPI enrichment p-value is shown for connected nodes and is considered significant if p<0.05 with individual PPI interaction significance not shown.

### 3.7 **Clusterin downregulates SMAD3, Tensin 3 (TNS3), GNAQ, and STAT1 transcriptionally.**

The important question posed was if the downregulation of various proteins related to regulators of actin dynamics was only at the protein level or was regulated at the transcript level. To address this, we performed qPCR-based gene expression analysis to evaluate the transcriptional impact of clusterin induction for selected downregulated proteins based on proteomics data. As illustrated in **Figure 6A**, clusterin gene expression was significantly induced following AdCLU treatment compared to AdMT (p=0.01, n=3). We found a significant decrease of SMAD3 (p=0.01, n=3) involved in TGFβ2 signaling pathway, TNS3 (p=0.03, n=3), GNAQ (p=0.04, n=3), and STAT1 (p=0.02, n=3). Transcripts of TNS1, TNS4, GNG12, RRAS2, STAT3, and STAT5b did not change. These results suggest that clusterin controls transcriptional regulation as well as at the protein level or only at the protein levels. Some of the proteins like GNG12 and RRAS2 are post-translationally modified via lipidation suggesting a post-translational regulation as well.

**Figure 6:**
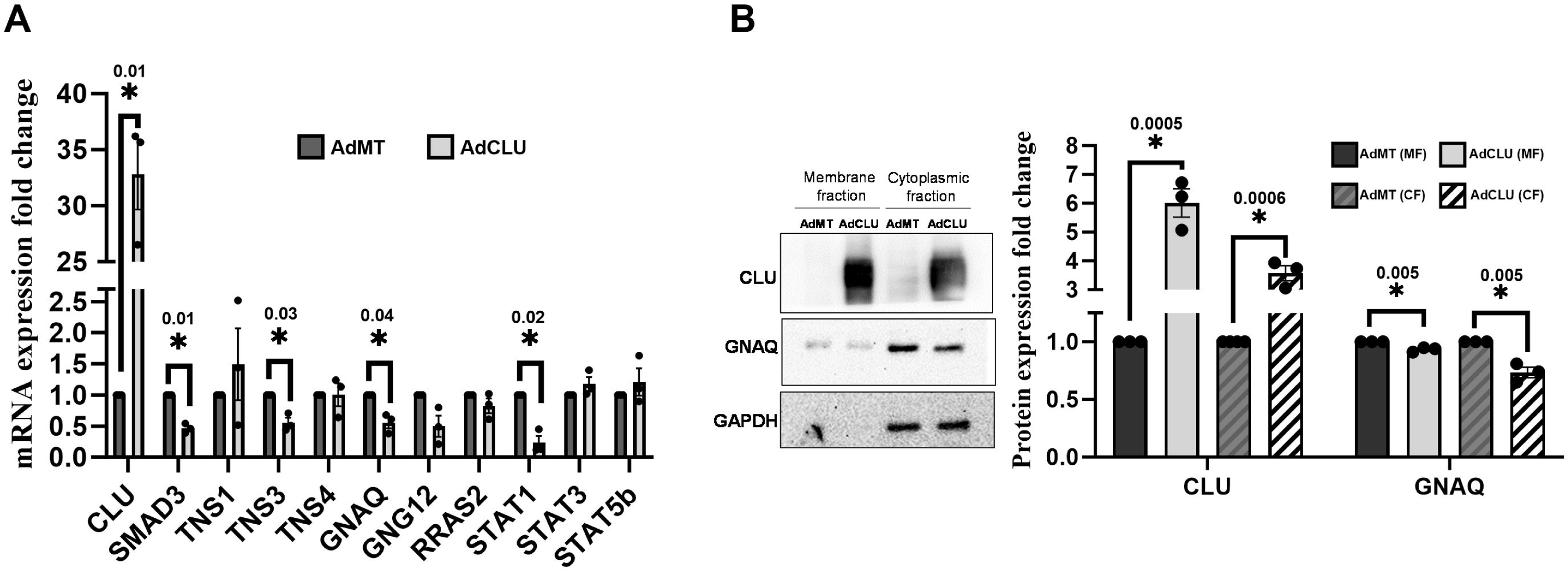
Transcriptional regulation of clusterin and effect of clusterin on GNAQ expression. A) Gene expression analysis showing the reduction in SMAD3, TNS3, GNAQ, and STAT1 under AdCLU compared to AdMT-treated HTM cells. B) IB showing membrane fraction (MF) and cytoplasmic fraction (CF) of PTM cells treated with either AdCLU or AdMT. AdCLU increased clusterin expression and reduced GNAQ expression in both MF and CF. Error bars represent the SEM, and (*) indicates significance, with a sample size of n=3.

### 3.8 **Clusterin reduces the localization of GNAQ to the membrane.**

GNAQ, integral to the core GTPase machinery, facilitates signal transmission from upstream GPCRs to downstream effectors such as PLCβ, initiating calcium signaling and protein kinase C (PKC) activation [9, 19, 50]. Although a decrease in gene expression was observed, this reduction did not translate to a noticeable decrease at the total protein level, but we found a significant reduction in ECM-enriched fraction. Considering the role of GNAQ as a membrane-associated protein that interacts with the inner surface of the plasma membrane, we aimed to analyze the effect of clusterin on GNAQ expression in both the membrane fraction (MF) and cytoplasmic fractions (CF) in PTM cells treated with AdCLU. As shown in **Figure 6B**, a significant increase in clusterin was observed under AdCLU treatment in both MF and CF (p=0.0005 and p=0.0006, respectively). GNAQ expression was significantly reduced in both MF (p=0.005) and CF (p=0.005) following clusterin induction. Notably, the reduction of GNAQ in the membrane fraction was smaller (10%) compared to the cytoplasmic fraction. This observation may suggest a relatively lower baseline availability of GNAQ at the membrane, or it could reflect the complex dynamics of GNAQ redistribution between the membrane and cytoplasm. The loss of GNAQ in the membrane fractions is potentially due to a decrease in post-translational modification in the form of lipidation (s-palmitoylation) in response to clusterin induction.

## 4. **Discussion**

This study revealed, for the first time, that clusterin plays a crucial role in mitigating pathological IOP elevation by attenuating the fibrotic changes in the TM outflow pathway. Mechanistically, clusterin negatively regulated the TM contractility and cell adhesive interactions by decreasing the cross-talk between acto-myosin networks and focal adhesions. Through comprehensive proteomic analyses, the study uncovered the significant impact of clusterin on TM function by differentially regulating the expression of proteins involved in cytoskeletal architecture, TGFβ2 signaling, protein folding, trafficking, and cellular quality control. Such multi-pronged targeting of key pathways involved in IOP regulation by clusterin implies a promising therapeutic avenue in treating ocular hypertension and glaucoma. Moreover, this study strongly suggests that clusterin is a critical protein in the TM and aqueous humor proteome involved in maintaining the IOP homeostasis.

Comprehensive analysis of the proteomic data unveiled the complex action of clusterin impacting biomechanical integrity of TM cells and essential signaling pathways like TGFβ2 and G-proteins. Of particular interest was the role of the clusterin in modulating the TGFβ2 signaling pathway, crucial for IOP regulation [30, 31, 51]. Our findings point to a marked inhibition of the canonical TGFβ2 signaling pathway by clusterin, as evidenced by the significant reduction in SMAD3 levels, a critical mediator of TGFβ2 signaling [14, 52]. This was consistent with our prior findings in mitigating TGFβ2-induced fibrotic responses [7]. Additionally, we had shown earlier that induction of TGFβ2 lowered the availability of functional clusterin by potentially inducing lysosomal degradation. The current study demonstrated that clusterin not only counteracts the IOP increases induced by TGFβ2 but also exerts a regulatory effect at the transcriptional level by significantly decreasing SMAD3 expression. Thus, suggesting that clusterin and TGFβ2 are under a perpetual loop to modulate each other’s bioavailability and maintain homeostasis. The transcriptional regulatory mechanism of TGFβ2 signaling by clusterin needs further investigation. Put together, these findings imply that a loss of clusterin can augment the effects of TGFβ2-mediated fibrogenic effects and increasing clusterin has the potential for IOP management.

Our study has uncovered a serendipitous and pivotal role of clusterin in the remodeling of the actin cytoskeleton within TM cells, a key determinant of aqueous humor drainage and IOP [9, 10, 12, 20]. The significant downregulation of actin-associated proteins such as ACTA1, adseverin, RHOQ, ACTN4, and Leiomodin-1 upon clusterin induction suggests a shift of TM towards a more relaxed state. This structural modification could improve aqueous humor drainage aiding in IOP reduction. Interestingly, the proteomics study found a significant lowering of actin and actin-related proteins. The ECM enrichment data showed minimal effect on ECM though the immunofluorescence studies showed greater impact on ECM. This could be a result of modifications in ECM secretion and cross-linking rather than ECM synthesis *per se*, which would not be reflected in the ECM enrichment.

Additionally, the remodeling of the actin cytoskeleton by clusterin affects not only the internal architecture of TM cells but also their cell-cell interactions and the interaction with ECM. The observed downregulation of proteins, such as tight junction protein ZO-1, filamin-binding LIM protein 1, and tensins, highlights how clusterin influences the communication between TM cells and the ECM. Specifically, the reduction in tensins - TNS1, TNS3, and TNS4 - which act as a liaison between the actin cytoskeleton and the ECM via focal adhesions is critical for cell matrix interactions. TNS1 and TNS3 are particularly interesting because they possess actin-binding domain (ABD)[53, 54]. The presence of actin-binding domains in TNS1 and TNS3 can link the cytoskeleton to the ECM [54], and exhibit changes that imply modifications in cell-ECM adhesion dynamics and possible alterations in cellular mechanotransduction. Particularly, TNS3 plays a critical role in the formation of fibrillar adhesion essential for fibronectin fibrillogenesis [55]. TNS3 knockout reduced fibrillar adhesions and fibronectin fibrillogenesis[55, 56]. The observed clusterin-mediated downregulation of TNS1 and TNS3 will have a strong impact on the cytoskeleton-ECM interactions and TM biomechanics. We propose this to be one of the reasons for the loss of focal adhesions and their maturation. Alterations in proteins like drebrin and thymosin beta-10, which play roles in regulating actin-myosin interactions [57] and actin sequestration [58], further corroborate to influence cellular contraction and adhesion, thereby impacting tissue stiffness and subsequently, IOP.

Interestingly, AdCLU was more effective than rhCLU at reducing actin polymerization and increasing TM contractility. This could be due to two possible mechanisms: 1) a constitutive increase in cellular clusterin, which may play an unknown role in remodeling actin and cell adhesion, and 2) the signaling of constitutively secreted clusterin through membrane-bound receptors compared to a set amount of rhCLU available for action. Future studies on these details should help understand the biology of clusterin in lowering IOP. Very little is known about the functional role of cellular clusterin, and it is an exciting prospect to better understand the functional significance of cellular clusterin. Studies on the role of clusterin on the impact of G protein signaling including - GNAQ, GNG12, GNB1, and RRAS2 – will be pivotal in targeting the downstream effectors, including PLCβ, and PKC in modulating actin dynamics, essential for maintaining the structural and functional integrity of the TM [9, 19, 59]. Additionally, our study also showed that clusterin significantly influenced the JAK-STAT signaling pathway. This was marked by the downregulation of key STAT family members like STAT1, STAT2, STAT3, and STAT5B. Downregulation of STAT along with the SMAD3 suggests concerted action of clusterin in reducing inflammatory and fibrotic responses in TM [60–62]. Additionally, the increase in proteins involved in ubiquitination pathways, notably UBE2R2 and UBE2G2 [63], suggests a possible pathway through which clusterin could promote the downregulation of ECM components and cytoskeletal proteins through ubiquitin-mediated degradation. The detailed mechanisms underlying this process remain to be elucidated in future research.

## 5. **Limitations and future directions**

While our study presents compelling evidence for the role of clusterin in IOP regulation, it acknowledges limitations. The *ex vivo* models, though insightful, may not fully replicate the complex dynamics of aqueous humor flow *in vivo*. Future studies should aim to validate these findings *in vivo*, exploring the long-term effects of clusterin modulation on IOP. Additionally, investigating the interaction of clusterin with other molecular players in the TM could unveil further therapeutic targets.

## Conclusion

In conclusion, our study demonstrates the pivotal role of clusterin in regulating IOP through intricate molecular mechanisms involving the actin cytoskeleton, ECM remodeling, and key signaling pathways. These insights not only enhance our understanding of clusterin but also open new avenues for therapeutic intervention for ocular hypertension and glaucoma.

## Acknowledgments

We would like to acknowledge Dr. Emma Doud from the Center for Proteome Analysis, IUSM, for helping with LC-MS/MS, data analysis, data evaluation, verification, and methods write-up. We thank Daniel Minner from Integrated Nanosystems Development Institute, Indiana University, for helping with Scanning Electron Microscopy imaging and analysis. This project was supported by the National Institutes of Health/National Eye Institute (R01EY029320) (PPP); Shaffer Grant from The Glaucoma Foundation (PPP); Award from the Ralph W. and Grace M. Showalter Research Trust and the Indiana University School of Medicine (PPP), Research Support Funds Grant (PPP), Cohen AMD Research Pilot Grant (PPP), RPB Departmental Pilot Grant (PPP), Glick Research Endowment Funds (PPP), and Challenge grant from Research to Prevent Blindness to IU; Postdoctoral Challenge Grant Indiana CTSI Core (EPAR1896) (AS).

## Data availability statement

The datasets presented in this study can be found in online repositories. The names of the repository and accession number(s) can be found in the article upon peer review.

## Author contributions

Conceptualization: PPP. Methodology: Cell cultures, immunofluorescence, immunoblotting, qPCR, and SEM: PPP and TW. Formal analysis: AS, PPP. Investigation: AS and PPP. Data curation: AS and PPP. Writing—original draft preparation: AS and PPP. Figure preparation: AS and TW. Writing—review and editing: AS and PPP. Visualization: AS and PPP. Supervision: PPP. Project administration: PPP. Funding acquisition: AS and PPP. All authors have read and agreed to the published version of the manuscript.

## Conflicts of Interest

The authors declare no conflicts of interest. The funders had no role in the design of the study; in the collection, analyses, or interpretation of data; in the writing of the manuscript, or in the decision to publish the results.

## Institutional Review Board statement

Ethical review and approval were waived for the use of cadaveric human eyes for *ex vivo* perfusion and the isolation of the TM cells from human cadaveric corneal rims for this study.

---

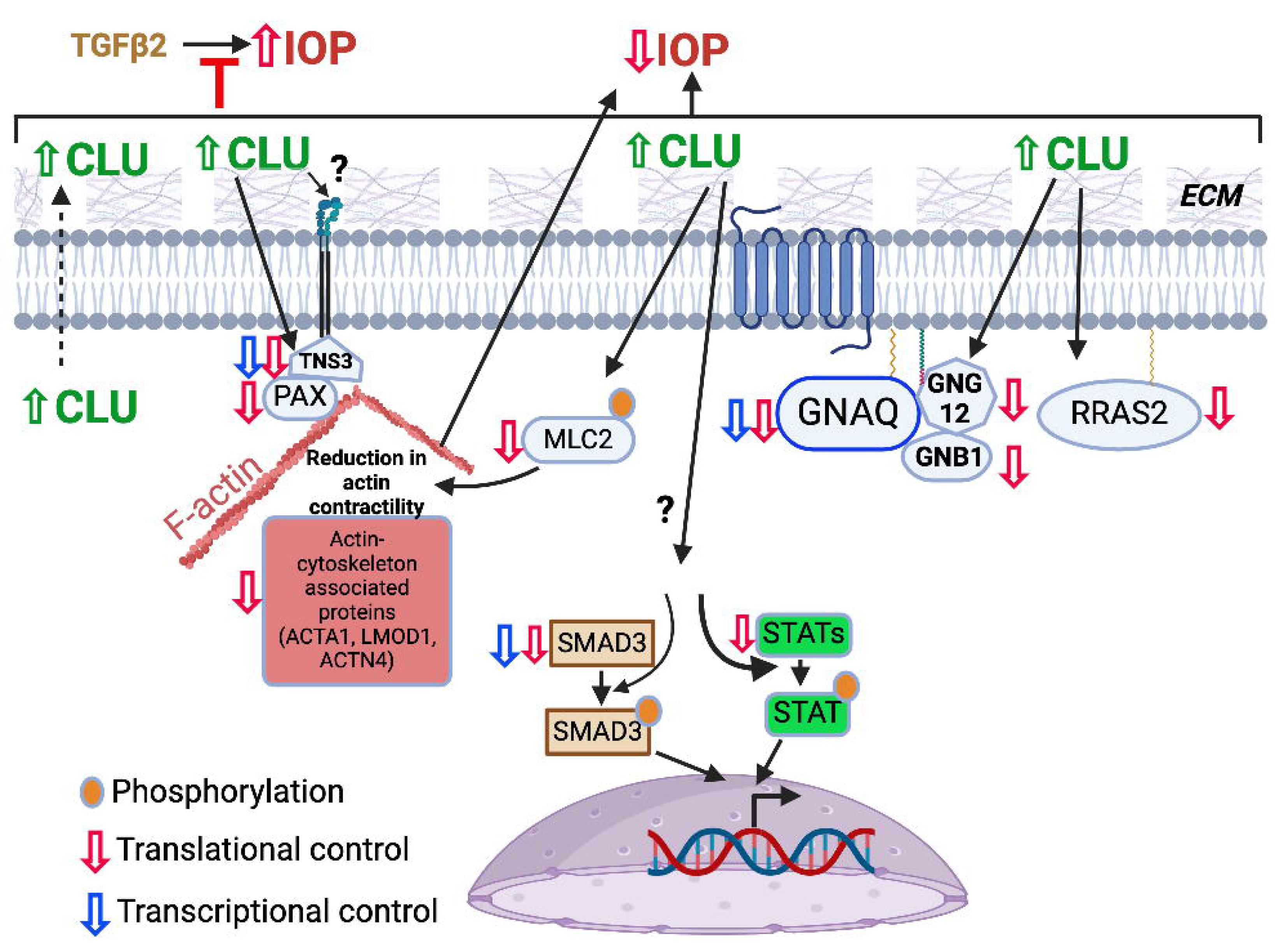

## Notes

### Competing Interest Statement

The authors have declared no competing interest.

## References

1. Tham, Y.C., et al., Global prevalence of glaucoma and projections of glaucoma burden through 2040: a systematic review and meta-analysis. Ophthalmology, 2014. 121(11): p. 2081–90.

2. Davis, B.M., et al., Glaucoma: the retina and beyond. Acta Neuropathol, 2016. 132(6): p. 807–826.

3. Quigley, H.A. and A.T. Broman, The number of people with glaucoma worldwide in 2010 and 2020. Br J Ophthalmol, 2006. 90(3): p. 262–7.

4. Sharif, N.A., Therapeutic Drugs and Devices for Tackling Ocular Hypertension and Glaucoma, and Need for Neuroprotection and Cytoprotective Therapies. Front Pharmacol, 2021. 12: p. 729249.

5. Goel, M., et al., Aqueous humor dynamics: a review. Open Ophthalmol J, 2010. 4: p. 52–9.

6. Pattabiraman, P.P. and C.B. Toris, The exit strategy: Pharmacological modulation of extracellular matrix production and deposition for better aqueous humor drainage. Eur J Pharmacol, 2016. 787: p. 32–42.

7. Soundararajan, A., et al., Novel insight into the role of clusterin on intraocular pressure regulation by modifying actin polymerization and extracellular matrix remodeling in the trabecular meshwork. J Cell Physiol, 2022. 237(7): p. 3012–3029.

8. El-Darzi, N., et al., The normalizing effects of the CYP46A1 activator efavirenz on retinal sterol levels and risk factors for glaucoma in Apoj(-/-) mice. Cell Mol Life Sci, 2023. 80(7): p. 194.

9. Pattabiraman, P.P., T. Inoue, and P.V. Rao, Elevated intraocular pressure induces Rho GTPase mediated contractile signaling in the trabecular meshwork. Exp Eye Res, 2015. 136: p. 29–33.

10. Pattabiraman, P.P., R. Maddala, and P.V. Rao, Regulation of plasticity and fibrogenic activity of trabecular meshwork cells by Rho GTPase signaling. J Cell Physiol, 2014. 229(7): p. 927–42.

11. Pattabiraman, P.P., et al., RhoA GTPase-induced ocular hypertension in a rodent model is associated with increased fibrogenic activity in the trabecular meshwork. Am J Pathol, 2015. 185(2): p. 496–512.

12. Rao, P.V., et al., Expression of dominant negative Rho-binding domain of Rho-kinase in organ cultured human eye anterior segments increases aqueous humor outflow. Mol Vis, 2005. 11: p. 288–97.

13. Patil, S.V., et al., A Novel Mouse Model of TGFβ2-Induced Ocular Hypertension Using Lentiviral Gene Delivery. Int J Mol Sci, 2022. 23(13).

14. McDowell, C.M., et al., Smad3 is necessary for transforming growth factor-beta2 induced ocular hypertension in mice. Exp Eye Res, 2013. 116: p. 419–23.

15. Rayana, N.P., et al., Using CRISPR Interference as a Therapeutic Approach to Treat TGFbeta2- Induced Ocular Hypertension and Glaucoma. Invest Ophthalmol Vis Sci, 2021. 62(12): p. 7.

16. Tovar-Vidales, T., A.F. Clark, and R.J. Wordinger, Transforming growth factor-beta2 utilizes the canonical Smad-signaling pathway to regulate tissue transglutaminase expression in human trabecular meshwork cells. Exp Eye Res, 2011. 93(4): p. 442–51.

17. Zhavoronkov, A., et al., Pro-fibrotic pathway activation in trabecular meshwork and lamina cribrosa is the main driving force of glaucoma. Cell Cycle, 2016. 15(12): p. 1643–52.

18. Hill, L.J., et al., TGF-beta-induced IOP elevations are mediated by RhoA in the early but not the late fibrotic phase of open angle glaucoma. Mol Vis, 2018. 24: p. 712–726.

19. Thieme, H., et al., The effects of protein kinase C on trabecular meshwork and ciliary muscle contractility. Invest Ophthalmol Vis Sci, 1999. 40(13): p. 3254–61.

20. Pattabiraman, P.P. and P.V. Rao, Mechanistic basis of Rho GTPase-induced extracellular matrix synthesis in trabecular meshwork cells. Am J Physiol Cell Physiol, 2010. 298(3): p. C749–63.

21. Yan, J., et al., Integrative transcriptomic and proteomic analysis reveals CD9/ITGA4/PI3K-Akt axis mediates trabecular meshwork cell apoptosis in human glaucoma. J Cell Mol Med, 2020. 24(1): p. 814–829.

22. Igarashi, N., M. Honjo, and M. Aihara, mTOR inhibitors potentially reduce TGF-β2-induced fibrogenic changes in trabecular meshwork cells. Scientific Reports, 2021. 11(1): p. 14111.

23. Yao, K., et al., Involvement of PI3K/Akt pathway in TGF-beta2-mediated epithelial mesenchymal transition in human lens epithelial cells. Ophthalmic Res, 2008. 40(2): p. 69–76.

24. Sabbineni, H., A. Verma, and P.R. Somanath, Isoform-specific effects of transforming growth factor beta on endothelial-to-mesenchymal transition. J Cell Physiol, 2018. 233(11): p. 8418–8428.

25. Rohne, P., et al., The chaperone activity of clusterin is dependent on glycosylation and redox environment. Cell Physiol Biochem, 2014. 34(5): p. 1626–39.

26. Keller, K.E., et al., Consensus recommendations for trabecular meshwork cell isolation, characterization and culture. Exp Eye Res, 2018. 171: p. 164–173.

27. Wang, T., et al., Identification of the novel role of sterol regulatory element binding proteins (SREBPs) in mechanotransduction and intraocular pressure regulation. FASEB J, 2023. 37(11): p. e23248.

28. Maddala, R. and V.P. Rao, *alpha-Crystallin localizes to the leading edges of migrating lens epithelial cells*. Exp Cell Res, 2005. 306(1): p. 203–15.

29. Dota, A., et al., Clusterin in human corneal endothelium and aqueous humor. Exp Eye Res, 1999. 69(6): p. 705–8.

30. Shepard, A.R., et al., Adenoviral gene transfer of active human transforming growth factor-beta2 elevates intraocular pressure and reduces outflow facility in rodent eyes. Invest Ophthalmol Vis Sci, 2010. 51(4): p. 2067–76.

31. Fleenor, D.L., et al., TGFbeta2-induced changes in human trabecular meshwork: implications for intraocular pressure. Invest Ophthalmol Vis Sci, 2006. 47(1): p. 226–34.

32. Alpha, K.M., W. Xu, and C.E. Turner, Paxillin family of focal adhesion adaptor proteins and regulation of cancer cell invasion. Int Rev Cell Mol Biol, 2020. 355: p. 1–52.

33. Busby, T., et al., Baf45a Mediated Chromatin Remodeling Promotes Transcriptional Activation for Osteogenesis and Odontogenesis. Front Endocrinol (Lausanne), 2021. 12: p. 763392.

34. Hodges, C., J.G. Kirkland, and G.R. Crabtree, The Many Roles of BAF (mSWI/SNF) and PBAF Complexes in Cancer. Cold Spring Harb Perspect Med, 2016. 6(8).

35. Orang, A., et al., Basonuclin-2 regulates extracellular matrix production and degradation. Life Sci Alliance, 2023. 6(10).

36. Tada, Y., et al., Ectonucleoside triphosphate diphosphohydrolase 6 expression in testis and testicular cancer and its implication in cisplatin resistance. Oncol Rep, 2011. 26(1): p. 161–7.

37. Bridges, R.J., N.R. Natale, and S.A. Patel, System xc(-) cystine/glutamate antiporter: an update on molecular pharmacology and roles within the CNS. Br J Pharmacol, 2012. 165(1): p. 20–34.

38. Mackenzie, B. and J.D. Erickson, Sodium-coupled neutral amino acid (System N/A) transporters of the SLC38 gene family. Pflugers Arch, 2004. 447(5): p. 784–95.

39. Yu, F.X., et al., Thymosin beta 10 and thymosin beta 4 are both actin monomer sequestering proteins. J Biol Chem, 1993. 268(1): p. 502–9.

40. Hayashi, K., et al., Modulatory role of drebrin on the cytoskeleton within dendritic spines in the rat cerebral cortex. J Neurosci, 1996. 16(22): p. 7161–70.

41. Winter, M., et al., Transient Receptor Potential Vanilloid 6 (TRPV6) Proteins Control the Extracellular Matrix Structure of the Placental Labyrinth. Int J Mol Sci, 2020. 21(24).

42. Ge, S.X., D. Jung, and R. Yao, ShinyGO: a graphical gene-set enrichment tool for animals and plants. Bioinformatics, 2020. 36(8): p. 2628–2629.

43. Millet, C. and Y.E. Zhang, Roles of Smad3 in TGF-beta signaling during carcinogenesis. Crit Rev Eukaryot Gene Expr, 2007. 17(4): p. 281–93.

44. Ropero, S., et al., Epigenetic loss of the familial tumor-suppressor gene exostosin-1 (EXT1) disrupts heparan sulfate synthesis in cancer cells. Hum Mol Genet, 2004. 13(22): p. 2753–65.

45. Qi, Y. and R. Xu, Roles of PLODs in Collagen Synthesis and Cancer Progression. Front Cell Dev Biol, 2018. 6: p. 66.

46. Bai, S.W., et al., Identification and characterization of a set of conserved and new regulators of cytoskeletal organization, cell morphology and migration. BMC Biol, 2011. 9: p. 54.

47. Gimm, J.A., et al., Functional characterization of spectrin-actin-binding domains in 4.1 family of proteins. Biochemistry, 2002. 41(23): p. 7275–82.

48. Ma, D., et al., *alpha-/gamma-Taxilin are required for centriolar subdistal appendage assembly and microtubule organization*. Elife, 2022. 11.

49. Lehtonen, S., F. Zhao, and E. Lehtonen, CD2-associated protein directly interacts with the actin cytoskeleton. Am J Physiol Renal Physiol, 2002. 283(4): p. F734–43.

50. Kostenis, E., E.M. Pfeil, and S. Annala, Heterotrimeric G(q) proteins as therapeutic targets? J Biol Chem, 2020. 295(16): p. 5206–5215.

51. Kasetti, R.B., et al., Transforming growth factor beta2 (TGFbeta2) signaling plays a key role in glucocorticoid-induced ocular hypertension. J Biol Chem, 2018. 293(25): p. 9854–9868.

52. Shen, X., et al., TGF-beta-induced phosphorylation of Smad3 regulates its interaction with coactivator p300/CREB-binding protein. Mol Biol Cell, 1998. 9(12): p. 3309–19.

53. Touaitahuata, H., et al., Tensin 3 is a new partner of Dock5 that controls osteoclast podosome organization and activity. J Cell Sci, 2016. 129(18): p. 3449–61.

54. Liao, Y.C. and S.H. Lo, Tensins - emerging insights into their domain functions, biological roles and disease relevance. J Cell Sci, 2021. 134(4).

55. Atherton, P., et al., Tensin3 interaction with talin drives the formation of fibronectin-associated fibrillar adhesions. J Cell Biol, 2022. 221(10).

56. Park, G.C., et al., Tensin-3 Regulates Integrin-Mediated Proliferation and Differentiation of Tonsil-Derived Mesenchymal Stem Cells. Cells, 2019. 9(1).

57. Ishikawa, R., et al., Drebrin attenuates the interaction between actin and myosin-V. Biochem Biophys Res Commun, 2007. 359(2): p. 398–401.

58. Zhang, X., et al., Thymosin beta 10 is a key regulator of tumorigenesis and metastasis and a novel serum marker in breast cancer. Breast Cancer Res, 2017. 19(1): p. 15.

59. Vazquez-Victorio, G., et al., GPCRs and actin-cytoskeleton dynamics. Methods Cell Biol, 2016. 132: p. 165–88.

60. Pedroza, M., et al., STAT-3 contributes to pulmonary fibrosis through epithelial injury and fibroblast-myofibroblast differentiation. FASEB J, 2016. 30(1): p. 129–40.

61. Banerjee, S., et al., JAK-STAT Signaling as a Target for Inflammatory and Autoimmune Diseases: Current and Future Prospects. Drugs, 2017. 77(5): p. 521–546.

62. Itoh, Y., M. Saitoh, and K. Miyazawa, Smad3-STAT3 crosstalk in pathophysiological contexts. Acta Biochim Biophys Sin (Shanghai), 2018. 50(1): p. 82–90.

63. Sharma, A., et al., Pharmacological Modulation of Ubiquitin-Proteasome Pathways in Oncogenic Signaling. Int J Mol Sci, 2021. 22(21).

