## Supplementary Tables for "Proteomic analysis uncovers clusterin-mediated disruption of actin-based contractile machinery in the trabecular meshwork to lower intraocular pressure"

**Supplementary Table 1: Details of donor age, sex, and race**

| **Experiment** | **Treatment** | **Age** | **Sex** | **Race** | **Eye details** |
| --- | --- | --- | --- | --- | --- |
| AdCLU perfusion study (n=5) | AdMT | 66 | Male | Caucasian | OS |
|  | AdMT | 47 | Female | Caucasian | OS |
|  | AdMT | 64 | Male | Caucasian | OD |
|  | AdMT | 70 | Female | Caucasian | OS |
|  | AdMT | 67 | Male | Caucasian | OD |
|  | AdCLU | 66 | Male | Caucasian | OD |
|  | AdCLU | 47 | Female | Caucasian | OS |
|  | AdCLU | 64 | Male | Caucasian | OS |
|  | AdCLU | 70 | Female | Caucasian | OD |
|  | AdCLU | 67 | Male | Caucasian | OS |
| rhCLU perfusion study (n=4) | rhCLU | 63 | Male | Caucasian | OS |
|  | rhCLU | 63 | Male | Caucasian | OD |
|  | rhCLU | 71 | Female | Caucasian | OS |
|  | rhCLU | 71 | Female | Caucasian | OD |
| TGFβ2 + rhCLU perfusion study (n=4) | TGFβ2 + rhCLU | 53 | Male | Caucasian | OS |
|  | TGFβ2 + rhCLU | 60 | Female | Caucasian | OD |
|  | TGFβ2 + rhCLU | 64 | Male | Caucasian | OS |
|  | TGFβ2 + rhCLU | 64 | Male | Caucasian | OD |
| Cells from TM tissues used for proteomics (n=4) | AdMT & AdCLU | 49 | Female | Caucasian | OD |
|  | AdMT & AdCLU | 68 | Male | Caucasian | OS |
|  | AdMT & AdCLU | 59 | Male | Caucasian | OS |
|  | AdMT & AdCLU | 73 | Female | Caucasian | OD |

**Supplementary Table 2: Pathway enrichment analysis using Shiny GO based on cellular component for upregulated cellular proteins upon treatment with AdCLU**

| **High level GO category** | **Genes** |
| --- | --- |
| **Organelle membrane** | DPM1 HCCS SLC25A5 ERP44 HSPA5 MSMO1 SEC61A1 TIGAR FDFT1 GNPTG RAB18 HM13 FNDC3A RAB2A YKT6 SEC61B GOSR1 RAB5C LRRC59 TMEM33 GRPEL1 SNX25 SC5D COPZ1   GOLT1B HSPA9 TTC1 SEC24A SSR3 SPCS1 SPCS2 RPN2 TMEM214 CLU SSR1 MSTO1 ERGIC3 SEC61G STT3A MRPL44 UQCC2 LRPPRC USP8 RNF185 SEC11A SLC16A3 RPTOR SH3GL1 TMCO1 SRPRB SNCA CISD2 YIPF5 HIGD2A NDUFB11 NSDHL ERLIN2 MTDH DNAJB12 HSD17B12 ARFGAP2 NDUFC2 UQCRB SLC35B2 RER1 GPAT4 MAIP1 RPN1 SLC30A8 VCP HSP90B1 PDIA3 SLC3A2 STX18 SLC25A6 TMED10 ORMDL3 DHCR7 LCLAT1 RAB1B BNIP3 MAN1B1 IMPDH2 CALR SRPRA COPB2 TMED9 TMEM120A ENTPD6 FAR1 RPS26 MRPL53 TMX2 ARFGAP3 MRPL20 DDOST TMEM199 PAM16 CANX |
| **Nuclear outer membrane-endoplasmic  reticulum membrane network** | DPM1 ERP44 HSPA5 MSMO1 SEC61A1 FDFT1 RAB18 HM13 RAB2A SEC61B LRRC59 TMEM33 SC5D COPZ1 SEC24A SSR3 SPCS1 SPCS2 RPN2 TMEM214 SSR1 ERGIC3 SEC61G STT3A LRPPRC RNF185 SEC11A TMCO1 SRPRB SNCA CISD2 YIPF5 NSDHL ERLIN2 MTDH DNAJB12 HSD17B12 SLC35B2 GPAT4 RPN1 VCP HSP90B1 PDIA3 STX18 TMED10 ORMDL3 DHCR7 LCLAT1 RAB1B MAN1B1 CALR SRPRA COPB2 TMED9 RPS26 TMX2 DDOST TMEM199 CANX |
| **Endoplasmic reticulum membrane** | DPM1 ERP44 HSPA5 MSMO1 SEC61A1 FDFT1 RAB18 HM13 RAB2A SEC61B LRRC59 TMEM33 SC5D COPZ1 SEC24A SSR3 SPCS1 SPCS2 RPN2 TMEM214 SSR1 ERGIC3 SEC61G STT3A   RNF185 SEC11A TMCO1 SRPRB CISD2 YIPF5 NSDHL ERLIN2 MTDH DNAJB12 HSD17B12 SLC35B2 GPAT4 RPN1 VCP HSP90B1 PDIA3 STX18 TMED10 ORMDL3 DHCR7 LCLAT1 RAB1B MAN1B1 CALR SRPRA COPB2 TMED9 RPS26 TMX2 DDOST TMEM199 CANX |
| **Extracellular region** | ERP44 HSPA5 ATP1B3 GNPTG MAGED2 RAB2A RPS16 RPL28 DDX5 RAB5C SLC38A1 HSPA9 ACAT2 CLU RPL5 COL5A1 TPT1 RPS15A RPS2 RPS11 RPL11 EIF2A MANF SNCA  PDIA4 RPL26 RPS14 VCP HSP90B1 PDIA3 SLC3A2 MUC15 FASN SYAP1 GNG5 RAB1B MAN1B1 RPLP2 IMPDH2 CALR RPL35A SRPRA CRELD2 TMED9 P4HB ENTPD6 RPS26  RPS4X RPL10A RPL39 RPS9 HYOU1 CANX |
| **Extracellular space** | ERP44 HSPA5 ATP1B3 GNPTG RAB2A RPS16 RPL28 DDX5 RAB5C SLC38A1 HSPA9 ACAT2 CLU RPL5 COL5A1 TPT1 RPS15A RPS2 RPS11 RPL11 EIF2A MANF SNCA PDIA4 RPL26  RPS14 VCP HSP90B1 PDIA3 SLC3A2 FASN SYAP1 GNG5 RAB1B RPLP2 IMPDH2 CALR RPL35A SRPRA CRELD2 TMED9 P4HB ENTPD6 RPS26 RPS4X RPL10A RPL39 RPS9 HYOU1 CANX |
| **Extracellular organelle** | ERP44 HSPA5 ATP1B3 GNPTG RAB2A RPS16 RPL28 DDX5 RAB5C SLC38A1 HSPA9 ACAT2 CLU RPL5 TPT1 RPS15A RPS2 RPS11 RPL11 RPL26 RPS14 VCP HSP90B1 PDIA3 SLC3A2  FASN SYAP1 GNG5 RAB1B MAN1B1 RPLP2 IMPDH2 CALR RPL35A SRPRA TMED9 P4HB RPS26 RPS4X RPL10A RPS9 HYOU1 CANX |
| **Cell junction** | HSPA5 RPL6 RAB18 PALMD RPS16 RPL18A GSK3A RPL19 EIF4G2 HSPA9 DBN1 CLU RPL5 USP8 RPS2 SLC16A3 SH3GL1 RPS11 SNCA RPL7 MTDH ITGA5 RPL9 RPS14 VCP HSP90B1 PDIA3 SLC3A2 SYAP1 RPS21 BNIP3 RPLP2 RPS27 CALR TMED9 P4HB RPS4X RPL10A RPS9 HYOU1 CANX |
| **Ribonucleoprotein complex** | RPL6 EIF3L RPS16 RPL18A RPL28 RPL19 DDX5 RPL5 EIF3G RPS15A MRPL44 RPS6 LRPPRC G3BP2 RPS2 RPS11 RPL11 EIF2A RPL10 RPL7 RPL26 RPL29 RPL9 RPS14 RPL13 RPS21 RPLP2 RPS27 CALR RPL35A RPS26 RPS4X RPL10A RPL39 MRPL53 ABCF1 MRPL20 RPL17 RPS9 |
| **Membrane protein complex** | SLC25A5 SEC61A1 ATP1B3 HM13 YKT6 SEC61B GOSR1 GRPEL1 COPZ1 HSPA9 SEC24A SPCS1 SPCS2 RPN2 SEC61G STT3A SEC11A SRPRB NDUFB11 HSD17B12 NDUFC2 UQCRB   ITGA5 RPN1 VCP PDIA3 SLC3A2 STX18 SLC25A6 TMED10 ORMDL3 GNG5 CALR SRPRA COPB2 DDOST TMEM199 PAM16 |
| **Envelope** | HCCS SLC25A5 TIGAR CHCHD2 LRRC59 TMEM33 GRPEL1 HSPA9 CLU MSTO1 MRPL44 UQCC2 LRPPRC RNF185 SLC16A3 SNCA CISD2 HIGD2A NDUFB11 CETN2 MTDH DNAJB12   NDUFC2 UQCRB MAIP1 SLC25A6 DHCR7 BNIP3 CALR XPOT TMEM120A MRPL53 TMX2 ABCF1 MRPL20 PAM16 |
| **Synapse** | RPL6 RAB18 PALMD RPL18A GSK3A RPL19 DBN1 CLU RPL5 USP8 SLC16A3 SH3GL1 SNCA RPL7 RPS14 VCP SLC3A2 SYAP1 RPS21 BNIP3 RPS27 TMED9 RPS4X RPL10A CANX |
| **Anchoring junction** | HSPA5 RPL6 RPS16 RPL19 EIF4G2 HSPA9 DBN1 RPL5 RPS2 RPS11 RPL7 MTDH ITGA5 RPL9 RPS14 HSP90B1 PDIA3 RPLP2 CALR P4HB RPS4X RPL10A RPS9 HYOU1 |
| **Cell-substrate junction** | HSPA5 RPL6 RPS16 RPL19 HSPA9 RPL5 RPS2 RPS11 RPL7 ITGA5 RPL9 RPS14 HSP90B1 PDIA3 RPLP2 CALR P4HB RPS4X RPL10A RPS9 HYOU1 |
| **Cell projection** | ATP1B3 PALMD GSK3A YKT6 RPL28 EIF4G2 SLC38A1 DBN1 CLU SLC34A1 RPS6 USP8 RPTOR SH3GL1 SNCA CETN2 SLC7A11 ITGA5 SYAP1 BNIP3 P4HB |
| **Intrinsic component of organelle membrane** | HSPA5 SEC61A1 HM13 TMEM33 HSPA9 SPCS1 ERGIC3 TMCO1 DNAJB12 SLC35B2 RER1 PDIA3 DHCR7 BNIP3 CALR FAR1 CANX |
| **Cell surface** | ERP44 HSPA5 HM13 SLC38A1 CLU SLC34A1 SLC7A11 PDIA4 RER1 ITGA5 PDIA3 SLC3A2 CALR P4HB ENTPD6 |
| **Postsynapse** | RPL6 PALMD RPL18A GSK3A DBN1 USP8 SLC16A3 SH3GL1 SNCA RPL7 RPS14 SYAP1 BNIP3 RPS27 RPL10A |
| **Plasma membrane region** | ATP1B3 RAB18 SLC38A1 DBN1 SLC34A1 SLC16A3 MTDH SLC7A11 ITGA5 SLC3A2 SYAP1 |
| **Neuron to neuron synapse** | RPL6 RPL18A DBN1 USP8 SLC16A3 SH3GL1 RPL7 RPS14 BNIP3 RPS27 RPL10A |
| **Postsynaptic specialization** | RPL6 RPL18A DBN1 USP8 SLC16A3 SH3GL1 RPL7 RPS14 BNIP3 RPS27 RPL10A |
| **Supramolecular fiber** | NUDC STUB1 GSK3A DBN1 TMEM214 COL5A1 TPT1 LRPPRC SRPRB SNCA |
| **Outer membrane** | TIGAR HSPA9 MSTO1 LRPPRC RNF185 SNCA CISD2 DHCR7 BNIP3 |
| **Cell body** | GSK3A YKT6 RPL28 SLC38A1 RPS6 RPTOR SNCA SLC3A2 SYAP1 |
| **Apical part of cell** | ATP1B3 RAB18 SLC38A1 SLC34A1 SLC16A3 CETN2 MTDH SLC7A11 SLC3A2 |
| **Side of membrane** | HM13 SLC38A1 SYAP1 GNG5 CALR P4HB RPS26 CANX |
| **Polymeric cytoskeletal fiber** | NUDC GSK3A DBN1 TMEM214 TPT1 LRPPRC SRPRB |
| **Nucleoid** | SLC25A5 LRRC59 HSPA9 UQCC2 LRPPRC LONP1 |
| **Mitochondrial nucleoid** | SLC25A5 LRRC59 HSPA9 UQCC2 LRPPRC LONP1 |
| **External encapsulating structure** | CLU COL5A1 HSD17B12 HSP90B1 CALR |
| **Midbody** | HSPA5 NUDC USP8 JTB HSP90B1 |
| **Extracellular matrix** | CLU COL5A1 HSD17B12 HSP90B1 CALR |
| **Presynapse** | SH3GL1 SNCA SYAP1 TMED9 CANX |
| **Cell-cell junction** | EIF4G2 DBN1 MTDH ITGA5 |
| **Extrinsic component of membrane** | USP8 SYAP1 GNG5 PAM16 |
| **Site of polarized growth** | FRYL DBN1 SNCA SYAP1 |
| **Basal part of cell** | ATP1B3 SLC38A1 SLC16A3 SLC3A2 |
| **Respirasome** | HIGD2A NDUFB11 NDUFC2 UQCRB |
| **Glutamatergic synapse** | DBN1 USP8 SH3GL1 VCP |
| **Oxidoreductase complex** | NDUFB11 NDUFC2 UQCRB P4HB |
| **Organellar ribosome** | MRPL44 MRPL53 MRPL20 |
| **Chromatin** | ASF1A HOXD9 PHF10 |
| **Nuclear outer membrane** | LRPPRC SNCA DHCR7 |
| **Cell leading edge** | DBN1 ITGA5 P4HB |
| **Coated membrane** | COPZ1 SEC24A COPB2 |
| **Synaptic membrane** | DBN1 SLC16A3 SYAP1 |
| **Lumenal side of membrane** | HM13 CALR CANX |
| **Cilium** | ATP1B3 CETN2 |
| **Synaptic vesicle** | SNCA TMED9 |
| **Organelle membrane contact site** | TMX2 CANX |
| **Blood microparticle** | CLU EIF2A |
| **Cluster of actin-based cell projections** | SLC34A1 SLC7A11 |
| **Synaptic vesicle membrane** | SNCA |
| **Protein-lipid complex** | CLU |
| **Plasma lipoprotein particle** | CLU |
| **Excitatory synapse** | SLC16A3 |
| **Cell pole** | SLC3A2 |
| **Intrinsic component of postsynaptic  membrane** | SLC16A3 |
| **Postsynaptic cytosol** | DBN1 |

**Supplementary Table 3: Pathway enrichment analysis using Shiny GO based on cellular component for downregulated cellular proteins upon treatment with AdCLU**

| **High level GO category** | **Genes** |
| --- | --- |
| **Extracellular region** | FUCA2 SCIN PAFAH1B1 MVP EHD2 HEXB WDR1 SRI ACAT1 FSCN1 GNB1 EPB41L2 CAD CPNE3 MAPK1 ARSA LGMN SMS NUTF2 CPPED1 PFN1 RAB35 SOD2 SNX3 ARSB MACROH2A1 NIT2 PRKAR2A CAPZA1 CRYZ CD46 NRP2 RHOQ PEPD AHNAK APOL2 MYO1B TWSG1 SNX9 ASS1 RRAS2 GSTM3 NAPG ITM2B ABI1 STXBP1 PRCP GGH RDX UACA ANXA7 ESD SCRN2 IGFBP4 ACTA1 ACP1 DYNLT1 MSN GSN PLBD2  QDPR GPD1L WASF2 FSTL1 BDH2 HDHD2 GLOD4 PAFAH1B2 SEPTIN2 TRAPPC1 GLB1 BSG GNG12 ARHGAP1 RNPEP BASP1 PLEC PPA1 IST1 TNFAIP2 RAB11B CDNF ADH5 CD2AP HEXA HSPA1B GART ACTN4 CC2D1A |
| **Extracellular space** | FUCA2 SCIN PAFAH1B1 MVP EHD2 HEXB WDR1 SRI ACAT1 FSCN1 GNB1 EPB41L2 CAD CPNE3 ARSA LGMN SMS NUTF2 PFN1 RAB35 SOD2 SNX3 ARSB MACROH2A1 NIT2 PRKAR2A CAPZA1 CRYZ CD46 RHOQ PEPD AHNAK MYO1B TWSG1 SNX9 ASS1 RRAS2 GSTM3 NAPG ITM2B ABI1 STXBP1 PRCP GGH RDX UACA ANXA7 ESD SCRN2 IGFBP4 ACTA1 ACP1 MSN GSN PLBD2 QDPR GPD1L WASF2 FSTL1 BDH2 HDHD2 GLOD4 PAFAH1B2 SEPTIN2 GLB1 BSG GNG12 ARHGAP1 RNPEP BASP1 PLEC PPA1 IST1 TNFAIP2 RAB11B CDNF ADH5 CD2AP HEXA HSPA1B GART ACTN4 CC2D1A |
| **Extracellular organelle** | FUCA2 SCIN PAFAH1B1 MVP EHD2 HEXB WDR1 SRI ACAT1 FSCN1 GNB1 EPB41L2 CAD CPNE3 ARSA LGMN SMS NUTF2 PFN1 RAB35 SOD2 SNX3 ARSB MACROH2A1 NIT2 PRKAR2A   CAPZA1 CRYZ CD46 RHOQ PEPD AHNAK MYO1B SNX9 ASS1 RRAS2 GSTM3 NAPG ITM2B ABI1 STXBP1 PRCP GGH RDX UACA ANXA7 ESD SCRN2 ACTA1 ACP1 MSN GSN PLBD2  QDPR GPD1L WASF2 FSTL1 BDH2 HDHD2 GLOD4 PAFAH1B2 SEPTIN2 GLB1 BSG GNG12 ARHGAP1 RNPEP BASP1 PLEC PPA1 IST1 RAB11B ADH5 CD2AP HEXA HSPA1B GART ACTN4 CC2D1A |
| **Cell junction** | CD99 SCIN PAFAH1B1 RAI14 PKN2 WDR1 SRI FSCN1 GNB1 EPB41L2 CAD CPNE3 MAPK1 TRIOBP PFN1 CCND1 RAB35 PRKAR2A CD46 NRP2 PDLIM2 AHNAK HNRNPH2 PALLD AKAP12 TNS4 MPRIP RRAS2 NAPG LMO7 TNS3 ABI1 STXBP1 RDX AKT1 MSN GSN WASF2 FBLIM1 NEXN JCAD SEPTIN2 BCL2L1 BSG MYD88 BASP1 PLEC RAB11B PDLIM7 CD2AP HSPA1B TJP1 ACTN4 |
| **Cell projection** | SCIN PAFAH1B1 PPP5C MRI1 PKN2 WDR1 SRI FSCN1 GNB1 CAD MAPK1 TRIOBP ZC3H14 RAB35 PRKAR2A STAT1 NRP2 MYO1B PALLD SNX9 ASS1 GSTM3 ABI1 STXBP1 RDX AKT1   ACTA1 DYNLT1 MSN GSN QDPR WASF2 NEXN JCAD SEPTIN2 BSG RAPH1 BASP1 PDLIM7 CD2AP SNX2 TJP1 ACTN4 |
| **Organelle membrane** | PAFAH1B1 BID EHD2 SRI GNB1 CPNE3 NUTF2 CCND1 RAB35 SNX3 EFHD1 SUMO1 CD46 RHOQ AHNAK APOL2 MYO1B SNX6 SNX9 ASS1 RRAS2 NAPG ITM2B SCRN1 PRCP ANXA7   LMNA SMAD3 SEPTIN2 BCL2L1 BSG MYD88 ARHGAP1 LMNB2 RAB11B SNX2 |
| **Anchoring junction** | CD99 PKN2 WDR1 FSCN1 EPB41L2 CPNE3 MAPK1 TRIOBP PFN1 CCND1 PRKAR2A CD46 PDLIM2 AHNAK PALLD AKAP12 TNS4 MPRIP RRAS2 LMO7 TNS3 RDX AKT1 MSN GSN  WASF2 FBLIM1 NEXN JCAD BSG PLEC PDLIM7 CD2AP HSPA1B TJP1 ACTN4 |
| **Cell-substrate junction** | CD99 EPB41L2 CPNE3 MAPK1 TRIOBP PFN1 PRKAR2A CD46 AHNAK PALLD AKAP12 TNS4 MPRIP RRAS2 LMO7 TNS3 RDX MSN GSN FBLIM1 NEXN BSG PLEC PDLIM7 HSPA1B ACTN4 |
| **Synapse** | PAFAH1B1 SRI GNB1 CAD MAPK1 PFN1 RAB35 PRKAR2A NRP2 HNRNPH2 PALLD AKAP12 NAPG ABI1 STXBP1 AKT1 WASF2 SEPTIN2 BCL2L1 MYD88 RAB11B |
| **Supramolecular fiber** | PAFAH1B1 SRI RHOQ PDLIM2 CALD1 AHNAK EML2 MYO1B PALLD ACTA1 DYNLT1 LMNA NEXN LMOD1 GNG12 LMNB2 RMDN1 PAWR PLEC PDLIM7 CD2AP |
| **Plasma membrane region** | EHD2 PKN2 FSCN1 GNB1 MAPK1 RAB35 PRKAR2A NRP2 LMO7 STXBP1 RDX MSN WASF2 JCAD SEPTIN2 CAVIN3 BSG CAVIN1 TJP1 |
| **Cell leading edge** | PAFAH1B1 PKN2 FSCN1 PALLD SNX9 ABI1 RDX AKT1 ACTA1 DYNLT1 GSN WASF2 JCAD RAPH1 PDLIM7 CD2AP SNX2 |
| **Polymeric cytoskeletal fiber** | PAFAH1B1 RHOQ PDLIM2 EML2 MYO1B PALLD ACTA1 DYNLT1 LMNA LMOD1 GNG12 LMNB2 RMDN1 PAWR PLEC PDLIM7 CD2AP |
| **Cell-cell junction** | PKN2 WDR1 FSCN1 CCND1 PDLIM2 AHNAK LMO7 RDX AKT1 WASF2 NEXN JCAD PDLIM7 CD2AP TJP1 |
| **Envelope** | PAFAH1B1 MVP BID NUTF2 CCND1 EFHD1 SUMO1 ASS1 LMO7 SCRN1 LMNA SMAD3 BCL2L1 LMNB2 IST1 |
| **Chromatin** | MACROH2A1 STAT1 ANP32E PSIP1 SMAD3 STAT5B BASP1 PAWR IST1 |
| **Cell body** | PAFAH1B1 PPP5C GNB1 CAD MAPK1 ASS1 AKAP12 ACTA1 DYNLT1 |
| **Postsynapse** | SRI MAPK1 NRP2 HNRNPH2 ABI1 STXBP1 AKT1 MYD88 |
| **Cilium** | PAFAH1B1 GNB1 PRKAR2A GSTM3 AKT1 SEPTIN2 BSG |
| **Apical part of cell** | LGMN MYO1B LMO7 RDX MSN TJP1 |
| **Side of membrane** | GNB1 ARSA PTP4A1 SNX9 AKT1 ACP1 GNG12 |
| **Presynapse** | SRI CAD RAB35 STXBP1 SEPTIN2 BCL2L1 RAB11B |
| **Membrane protein complex** | GNB1 SNX3 SUMO1 SNX6 NAPG GNG12 SNX2 |
| **Extrinsic component of membrane** | EHD2 GNB1 ARSA SNX9 AKT1 GNG12 |
| **Site of polarized growth** | PAFAH1B1 FSCN1 PALLD ABI1 DYNLT1 BASP1 |
| **Midbody** | PKN2 TRIOBP EXOC2 RDX SEPTIN2 IST1 |
| **Ribonucleoprotein complex** | CIRBP ZC3H14 HNRNPH2 VBP1 HSPA1B ACTN4 |
| **Cell surface** | ARSA ARSB CD46 LMO7 MSN |
| **Nuclear outer membrane-endoplasmic reticulum membrane network** | SRI NUTF2 APOL2 ANXA7 BSG |
| **Basal part of cell** | PRCP MSN WASF2 BSG TJP1 |
| **Blood microparticle** | PFN1 ACTA1 MSN GSN HSPA1B |
| **Endoplasmic reticulum membrane** | SRI APOL2 ANXA7 BSG |
| **Synaptic vesicle** | RAB35 SEPTIN2 BCL2L1 RAB11B |
| **Outer membrane** | BID NUTF2 ASS1 BCL2L1 |
| **Intrinsic component of organelle membrane** | BID RAB35 ITM2B RAB11B |
| **Intercellular bridge** | EHD2 RAB35 GSTM3 SEPTIN2 |
| **Cluster of actin-based cell projections** | PAFAH1B1 TRIOBP MYO1B PLEC |
| **Neuron to neuron synapse** | MAPK1 HNRNPH2 ABI1 MYD88 |
| **Postsynaptic specialization** | MAPK1 HNRNPH2 ABI1 MYD88 |
| **Synaptic vesicle membrane** | RAB35 BCL2L1 RAB11B |
| **Retromer complex** | SNX3 SNX6 SNX2 |
| **Cell division site** | PKN2 RDX SEPTIN2 |
| **Glutamatergic synapse** | PFN1 NRP2 STXBP1 |
| **External encapsulating structure** | ANXA7 |
| **Retromer, tubulation complex** | SNX6 SNX2 |
| **Extracellular matrix** | ANXA7 |
| **Stereocilium** | PAFAH1B1 TRIOBP |
| **Flemming body** | EXOC2 IST1 |
| **Synaptic membrane** | NRP2 STXBP1 |
| **Intrinsic component of synaptic  vesicle membrane** | RAB35 RAB11B |
| **Nucleoid** | SOD2 |
| **Nuclear outer membrane** | NUTF2 |
| **Clathrin-coated pit** | RAB35 |
| **Host cellular component** | DYNLT1 |
| **Cell trailing edge** | MSN |
| **Protein-DNA complex** | MACROH2A1 |
| **Methyltransferase complex** | CLNS1A |
| **Mitochondrial nucleoid** | SOD2 |
| **Myelin sheath** | GSN |
| **Dendritic spine neck** | SRI |
| **Presynaptic active zone** | STXBP1 |
| **Cell tip** | RDX |
| **Excitatory synapse** | PALLD |
| **Cell pole** | RDX |
| **Intrinsic component of postsynaptic  membrane** | NRP2 |
| **Oxidoreductase complex** | GPD1L |
